# Negative legacy effects of past forest use on plant diversity in semi-natural grasslands on ski slopes

**DOI:** 10.1101/2022.08.11.503702

**Authors:** Yuki A. Yaida, Takuma Nagai, Kazuya Oguro, Koki R. Katsuhara, Kei Uchida, Tanaka Kenta, Atushi Ushimaru

**Author notes:** (Y.A. Yaida); (T. Nagai); (K. Oguro); (K.R. Katsuhara); (K. Uchida); (T. Kenta); (A. Ushimaru). YAY is a contact author and AU is a correspondence author.

## Abstract

Over the past century, grassland and forest ecosystems globally have been heavily influenced by land use changes driven by diverse socioeconomic activities. Ski resorts are a modern land-use type associated with biodiversity loss in mountain ecosystems worldwide. Below the treeline, by contrast, some ski slopes have been shown to provide suboptimal semi-natural habitats for native grassland plants and animals, depending on specific construction and management practices. We compared environmental factors and grassland vegetation between two types of ski slopes in central Japan with different land-use histories: slopes constructed on old pastures (pasture slopes) and slopes constructed by clearing secondary forests or *Larix kaempferi* plantations established on abandoned pastures during the 1940s–1990s (forest slopes). We examined the effects of land use history and machine grading as well as other environmental factors on ski slope vegetation, including total species richness and the richness of native, endangered, and exotic plants, using a total of 108 plots of 2 m × 10 m. Compared to pasture slopes, forest slopes exhibited significantly lower richness of total plants and native grassland species, including endangered species. Forest slopes were more graded than pasture slopes, resulting in lower native and higher exotic grassland species richness. A significantly lower duration of direct sunlight on forest slopes than on pasture slopes possibly decreased endangered species diversity. The lower species richness on forest slopes may be partly caused by seed dispersal limitations. Our findings demonstrate that ski slopes have good potential to support numerous native grassland plant species, including endangered species, but this potential is significantly and negatively affected by forest use history and concomitant environmental changes. The conservation of semi-natural conditions on pasture slopes as habitats for native grassland species can be promoted through the maintenance of annual mowing practices, avoidance of machine grading, and wider ski courses.

**Nomenclature:** Yonekura & Kajita (2003) BG Plants: index of Japanese names and scientific names (YList; http://ylist.info/index.html)

## Introduction

Land use change is considered the primary driver of global biodiversity decline over the past century (Sala et al., 2000; Gerstner, Dormann, Stein, Manceur, & Seppelt, 2014). The negative effects of land-use change on biodiversity, as well as the mechanisms underlying these effects, have been intensively studied in both natural and semi-natural ecosystems (Butchart et al., 2010; Newbold et al., 2015). However, the positive effects of land-use changes on biodiversity maintenance, conservation, and restoration have rarely been examined. In recent decades, grassland and forest ecosystems have been heavily impacted by land-use changes, such as monocultural plantations, intensive agriculture, land abandonment, and urbanization, resulting from changing global socioeconomic pressures (Foley et al., 2005; Uchida & Ushimaru, 2015; To□ro□k & Dengler, 2018). Although some reports have shown that modern land uses, such as power line corridors, golf courses, and ski slopes (Tsuyuzaki, 2002; Yasuda & Koike, 2006; Lampinen, Ruokolainen & Huhta, 2015) maintain threatened semi-natural grassland conditions, we did not find sufficient conservational perspectives for grassland specific species in such newly created habitats.

Ski resorts are a modern land use type associated with biodiversity losses in mountain ecosystems (Tsuyuzaki, 1994; Wipf, Rixen, Fischer, Schmid, & Stoeckli, 2005; Rolando, Caprio, Rinaldi, & Ellena, 2007; Burt & Rice, 2009; Roux-Fouillet, Wipf, & Rixen, 2011). The construction and maintenance of ski slopes degrades natural alpine grass-, and shrub-land (Wipf et al., 2005; Roland et al., 2007; Roux-Fouillet et al., 2011), as well as subalpine and montane forests (Tsuyuzaki, 1994; Burt & Rice, 2009; Roland et al., 2007). Damage results from the removal of trees and shrubs, machine grading (scraping and levelling off the ground surface with heavy machinery, Fig. S1a), artificial snowmaking, and the introduction of exotic species for erosion control (Fig. S1b). These practices have been demonstrated to have long-lasting negative effects on plant communities in ski resorts in North America, Europe, and East Asia (Tsuyuzaki, 1994; Wipf et al., 2005; Roland et al., 2007; Burt & Rice, 2009; Inoue et al., 2021). Clearing results in wholesale community conversion from forests or shrublands to grasslands, whereas grading and artificial snowmaking typically affect surface soil properties, such as the cover of bare ground and soil moisture and nutrient levels, which, in turn, affect plant diversity and composition (Tsuyuzaki, 1995, 2002; Roland et al., 2007; Burt & Rice, 2009; Roux-Fouillet et al., 2011). Soil erosion resulting from grading leads to reduced water content and soil depth (Watson, 1985; Tsuyuzaki, 1990), whereas the introduction of exotic vegetation after grading often promotes dominance of these plant species during the early stages of succession (Tsuyuzaki, 1995, 2002).

In contrast, some ski slopes in Europe, North America, and East Asia have been shown to provide suboptimal habitats for native grassland plants and animals below the treeline, depending on the specific construction and management practices (Burt & Rice, 2009; Roland et al., 2007; Tsuyuzaki, 1995). Most ski resorts in Japan have been constructed by clearing, grading, and planting exotic species in areas with well-developed forests and heavy snow cover (Tsuyuzaki, 1994). Despite their negative impacts on forest communities, ski slopes below the treeline can sustain semi-natural grassland habitats (Tsuyuzaki, 1995; Roland et al., 2007), which have been lost in recent decades in the Palearctic region (Ogura, 2006; To□ro□k & Dengler, 2018; Ushimaru, Uchida, & Suka, 2018; Ushimaru, Uchida, Ikegami & Suka, 2021). Moreover, some Japanese ski slopes have been established and maintained on old pastures without intensive grading. Recently, ski slopes on old pastures (pasture slopes) have been found to provide habitats for endangered grassland plants and butterflies (Nakahama, Uchida, Ushimaru, & Isagi, 2018; Yaida et al., 2019), which are currently threatened by land use changes occurring throughout semi-natural grassland areas in Japan (Koyanagi & Furukawa, 2013; Uchida, Takahashi, Shinohara, & Ushimaru, 2016).

Inoue et al. (2021) recently reported that ski slopes constructed by clearing forests established on abandoned pastures (forest slopes) have maintained poorer grassland plant richness than pasture slopes, although the current management practices were the same between the two types of ski slopes. To date, the reason forest slopes harbor fewer grassland species, including endangered species, compared to pasture slopes, has not been examined. Forest slopes largely lack endangered species, even if the slopes were created more than 50 years ago (Yaida et al., 2019; Inoue et al., 2021). To validate the potential of forest slopes for grassland habitat conservation, we should clarify the underlying mechanisms of the legacy effects of short-term forestation (for ca. 25–50 years) on grassland species richness by comparing both vegetation and environmental factors between pasture and forest slopes. This issue was not examined in our previous studies (Yaida et al., 2019; Inoue et al., 2021) and is worth studying even for the restoration of semi-natural grassland vegetation in forested areas.

In this study, we examined environmental factors and their relationships with vegetation components on ski slopes with two different land-use histories to elucidate the mechanisms of the legacy effects of past forestation on current grasslands. First, pasture slopes that were constructed on old pastures in 1928 and second, forest slopes constructed by clearing secondary forests or conifer plantations established on abandoned pastures during the 1940s–1990s (Fig. S1c). Based on old cadastral maps (Inoue et al., 2021), we confirmed that all the study slopes were used as pastures approximately 160 years ago. Thus, it is important to note that grassland communities on forest slopes have been reestablished after slope construction. We examined the effects of forested history and machine grading on the environmental and vegetation conditions of the ski slopes. Here, we hypothesized that the poorer grassland plant richness of forest slopes is caused by changes in soil and light conditions owing to forestation and heavy machine grading (the environmental change hypothesis). Moreover, given that grassland vegetation was removed by forestation (Inoue et al., 2021) and recovery on forest slopes would be reliant on seeds dispersed from neighboring grasslands (Öster, Ask, Cousins, & Eriksson, 2009), we can also propose another hypothesis that seed dispersal limitation has prohibited the re-establishment of some grassland plants, including endangered species, on forest slopes (the dispersal limitation hypothesis). The dispersal limitation hypothesis predicts that species with lower dispersal ability and/or lower abundance in species-rich source grasslands (pasture slopes in this study) are limited to forest slopes. We explored factors depressing the potential of forest slopes as habitats for threatened grassland plants to promote the restoration and conservation of grassland biodiversity.

## Materials and methods

### Study area, sites, and plots

Field data were collected in 2015 and 2016 on 25 ski slopes at seven ski resorts in Nagano and Gunma Prefectures, Japan (Fig. 1), seven slopes at Dabosu, six at Taro, one at Omatsu, two at Tsubakuro, and three each at the Minenohara, Yunomaru, and Kazawa ski resorts (Table S1). The mean annual temperatures in the study areas are 6.0–8.0°C. The mean coldest month (January) minimum temperatures range from −5.8 to −3.3°C, while the average maximum temperatures in the warmest month (July) are 19.1–19.4°C. The mean annual precipitation between 1986 and 2015 was 1197.3–1500 mm. The cumulative and maximum snowfall depths during winter to early spring (from November to April) between 1986 and 2015 were 294–831 cm and 59–152 cm, respectively. Meteorological data were collected by the Japanese Meteorological Agency (https://www.jma.go.jp/jma/indexe.html) at two nearby automated weather stations located at 36°31.9’ N, 138°19.5’ E (1253 m a.s.l.) and 36°27.8’ N, 138°27.8’ E (1230 m a.s.l.).

**Fig. 1.**
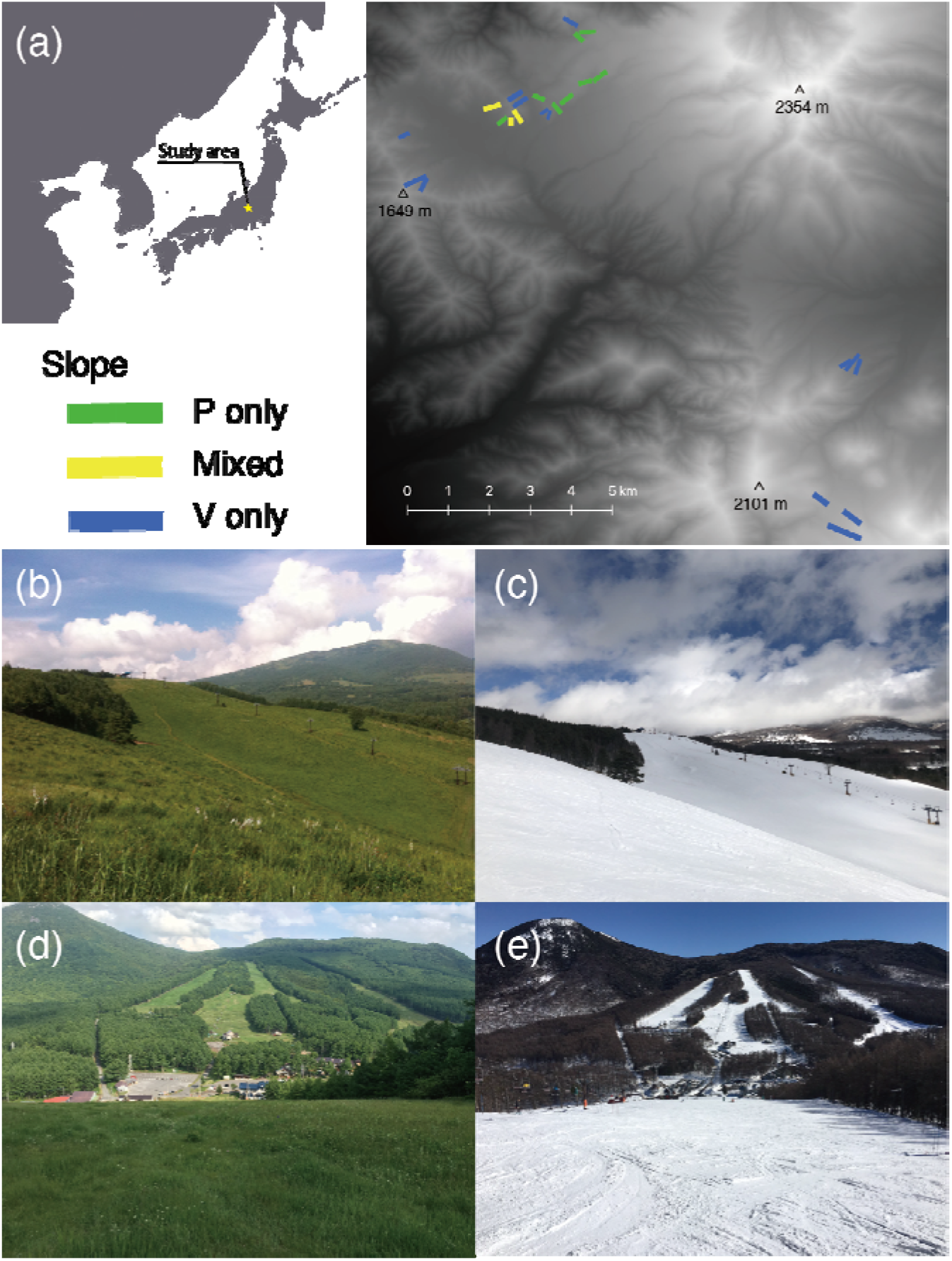
Location of the study area and distribution of study slopes (a). Green and blue lines indicate study slopes containing four pasture (P only) and four forest (F only) plots, respectively. Yellow lines represent the four slopes that include both pasture and forest plots (mixed). Photographs of pasture slopes in summer (b) and in winter (c), and forest ski slopes in summer (d) and in winter (e).

All study slopes were located in the cool temperate zone (1327–1882 m a.s.l.). Although the climax vegetation of the study area comprises broad-leaved deciduous forests, ski slopes are maintained by annual mowing in September and cut materials are left *in situ*, resulting in litter accumulation. One slope at the Yunomaru ski resort was used for extensive cattle grazing in the summer. Artificial snow-making was applied to all study slopes depending on snowfall conditions.

At each site, we established four 2 m × 10 m plots, located at the center and edge of the slope, and at the upper and lower slope positions (Fig. S2a). We surveyed totally 100 plots on slopes as well as an additional eight 2 m × 10 m plots in a semi-natural grassland at the Sugadaira Mountain Research Centre (SMRC, 36°31’24” N, 138°20’52” E, 1328 m a.s.l.) at Tsukuba University, which is 1 km away from the Dabosu slopes (Fig. S1d). This site has been maintained as a meadow with an annual autumn mowing regime for approximately 80 years following its historical use as pasture land, and the plots surveyed here served as reference grassland plots.

### Land use history

We determined the land use history of each plot using cadastral maps from 1910–12 and 1930–37, aerial photographs from 1948, 1965, 1975, 1991, 1996, and 2010, provided by the Geospatial Information Authority of Japan. We also referred to an old land-use map (*Sugadaira kaikon no zu*), which showed that pastures covered almost the entire study area in 1855. Thirty-three plots were on slopes that had been converted directly from pastures (pasture plots), where grassland conditions had been continuously maintained for at least 160 years (Table S1). The other 67 plots (forest plots) were located where secondary forests or conifer plantations were converted to ski slopes between 1912 and 1996. The three slopes included both pasture and forest plots.

### Vegetation surveys

We conducted vegetation surveys in late June and early August 2015 in 76 plots on 19 slopes, as well as in the eight reference plots at SMRC. In 2016, we surveyed an additional 24 plots on six slopes. We recorded all vascular plant species observed within each plot and pooled the data from the June and August surveys for the analysis. The total species richness, native grassland richness, and exotic species richness in each plot were used as diversity indices. We referred to the wild flowers of Japan (Satake et al. 1981) for the identification of grassland species. We estimated the original abundance of each grassland species by counting the number of pasture plots in which the species was recorded and categorized their dispersal modes into seven types: barochory (gravity), ballochory (explosive dehiscence), hydrochory (water, mostly raindrops), myrmecochory (ants), anemochory (wind), epizoochory (mammals), and endozoochory (mostly birds).

We also calculated the Red List Index value (RLI) for each plot based on the 47 Prefecture Red Lists in Japan as an indicator of endangered species diversity. We used a modification of the formulas developed by Butchart et al. (2010) to calculate the index (Eq. 1).

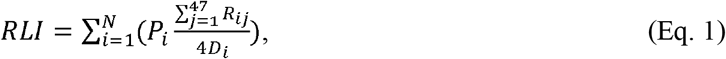

where *P_i_* is the presence/absence (1/0) of species *i* in a given plot and *R_ij_* is the Red List rank of species *i* in prefecture *j*, where 4 = extinct, 3 = critically endangered or endangered (CR and EN), 2 = vulnerable (VU), 1 = near-threatened (NT), and 0 = others. *D_i_* is the number of prefectures in which species *i* is distributed and *N* is the total number of plant species observed in this study (279).

Thus, we first calculated the average threatened status per prefecture of species *i* (Eq. 2).

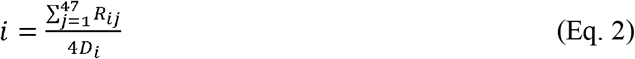

If a given species was VU in all distributing prefectures, the value was 0.5. We then summed the values of all species that occurred in the target plot. A high RLI value indicated a high diversity of species, with a declining trend throughout their distribution range in the plot.

Moreover, we examined the richness of endangered species, defined as species listed as NT or higher on the national Red List (Ministry of the Environment of Japan, 2019). Endangered species richness data of 47 out of 108 plots (mostly pasture plots) were published by Yaida et al. (2019).

### Vegetation and environmental factors

We recorded vegetation height, aboveground biomass, soil pH, the degree of soil grading, slope angle, and duration of direct sunlight during the summer solstice (hereafter, duration of direct sunlight) in early August for each plot on 19 slopes in 2015, and six in 2016. We measured the height (in cm) of the tallest plant within a 10 cm radius of each of the five points in the plot (Fig. S2b) and used the mean value to represent the vegetation height for analysis. To quantify aboveground biomass, we clipped all live aboveground vascular plant material from two additional 0.25 × 0.25 m^2^ subplots located adjacent to each plot. We dried the clipped material at 70°C for 72 h, weighed the dry biomass for each subplot, and averaged the values from the two subplots for use in the analysis.

Concurrently with the biomass sampling, we collected soil samples from the two subplots (see Fig. S2) at a depth of 0–5 cm, using a 5 × 5 cm cylindrical soil corer (approximately 100 cm^3^). The samples from each plot were combined and dried at 70°C for 72 h. Soil pH was measured for the mixed samples and dried soil samples in a 2:5 (w:w) soil:distilled water mixture using a HI 991221 Direct Soil pH measurement instrument (HANNA Instruments, Smithfield, RI, USA).

The Japanese black soil *Kurobokudo* (known as Andosols in the FAO Soil classification), is a volcanic soil rich in type A humic acid. It typically occurs in topsoil of old semi-natural grasslands that have been maintained over the long-term (Kawano, Sasaki, Hayashi, & Takahara, 2012; Miyabuchi, Sugiyama, & Nagaoka, 2012; Ushimaru et al., 2018). In the study area, *Kurobokudo* is widely distributed in topsoil to a depth of approximately 50 cm (Fig. S1e). Machine grading removed *Kurobokudo* and exposed the volcanic subsoil on the study slopes (Fig. S1a). To estimate the grading intensity, we assessed the Melanic Index (MI) of each soil sample. MI is an indicator that can easily distinguish *kurobokudo* from other volcanic soils and is widely used by Japanese pedologists (see detail Honna, Yamamoto & Matsui, 1988; Yamamoto et al., 2000). In Japan, *Kurobokudo* normally exhibits an MI ≤ 1.7, whereas volcanic subsoils have larger values of MI (Honna et al., 1988; Takahashi & Shoji, 2002); thus, we assume that MI increases with the intensity of grading. The slope angle of each plot was measured by using a clinometer. Steeper slopes have often been created for advanced skiers by using machine grading.

As an index of light conditions during the growing season, we examined the duration of direct sunlight using hemispherical photographs taken at 1 m above ground level in the center of each plot in overcast conditions in late August. We used the analysis program CanopOn 2 (takenaka-akio.org/etc./canopon2/ index.html; cf. Ohara and Ushimaru, 2015) to quantify the duration of direct sunlight.

Data of aboveground biomass, soil pH, slope angle, and the duration of direct sunlight of 47 out of the 108 plots were already reported in a previous study (Yaida et al., 2019).

### Statistical analyses

#### Effects of land use history and grading on vegetation and environmental conditions

We first constructed generalized linear models (GLMs) using a Gaussian distribution and an identity link to examine whether forest plots were more graded than pasture slopes. In the model, land-use history type (pasture/forest plot, 0/1) and slope angle were incorporated as explanatory variables and the intensity of grading (using MI as a proxy) response variables, respectively. Because the study slopes and plots were not randomly distributed, there was potential for spatial autocorrelation to affect the results. Therefore, we calculated the spatial autocovariate (a distance-weighted function of neighboring response values) for each plot using latitude and longitude and incorporated the values into this and all the following GLMs as a covariate (Dormann et al. 2007).

We then constructed GLMs (a Gaussian distribution and an identity-link function) to examine the effects of land-use history, MI, and slope angle on four vegetation and environmental factors (vegetation height, aboveground biomass, soil pH, and duration of direct sunlight). In the models, land use history type, MI, and autocovariates were incorporated as independent explanatory variables, while each of the four environmental factors was the response variable. Prior to the analyses, a variance inflation factor (VIF) test detected only weak multicollinearity among the explanatory variables in the full model (VIFs < 3). Therefore, we included all the variables in the models.

#### Species richness

We then constructed GLMs (Poisson distribution and log-link function) to examine the effects of land-use history, vegetation, and environmental factors on three plant diversity indices. In the models, land use history type, MI, vegetation height, aboveground biomass, soil pH, duration of direct sunlight, and slope angle were incorporated as explanatory variables, whereas total, native grassland, endangered grassland, and exotic species richness were the response variables. We constructed an additional GLM using a Gaussian distribution and a log-link function, in which RLI was incorporated as the response variable; the explanatory variables were the same as in the previous GLMs. Because the study slopes and plots were not randomly distributed, we incorporated the spatial autocovariate values into each GLM as a covariate (Dormann et al. 2007). We conducted a VIF test to confirm only weak multicollinearity among the explanatory variables in the full model (VIFs < 3).

We also constructed GLMs to examine the potential of ski slopes as semi-natural grasslands. In the models, land use history type (reference grassland at SMRC/pasture/forest) and the autocovariate were incorporated as explanatory variables, while three richness (Poisson distribution and log-link function) values and RLI (Gaussian distribution and log-link function) were the response variables. The distribution of eight endangered species on the national Red List and their relationships with vegetation and environmental variables in forty-seven (mostly pasture) plots out of the 108 study plots, have already been published (Yaida et al. 2019). In this study, we added data on total species, including common native and exotic plants, and those from 61 forest and reference plots to elucidate differences in environmental factors and vegetation between plot types.

#### Species composition

We used nonmetric multidimensional scaling (NMDS), which has been demonstrated to be robust and effective for community data (Minchin, 1987), to graphically examine differences in species composition between pasture and forest plots. We calculated among-plot dissimilarities using the Jaccard dissimilarity index (JDI) for all plot pairs using data for all species. We used a two-dimensional NMDS graph to visualize differences in species composition, and then fitted vectors for land-use history type (pasture/forest, 0/1), MI, vegetation height, aboveground biomass, soil pH, duration of direct sunlight, slope angle, and richness metrics, and examined the significance of each vector based on 10,000 random permutations using the *envfit* function in the R package *vegan*.

#### Effects of dispersal limitation

To examine the effects of dispersal limitation on the establishment of grassland species in forest plots, we examined the relationship between relative species abundance in pasture and forest plots using generalized linear mixed models (GLMMs) with a Gaussian distribution and an identity-link function. The abundance of each species in the forest plots, that is, the number of forest plots where the species occurred divided by the total number of forest plots, was the response variable. Explanatory variables included abundance in pasture plots, species group (0/1, native grassland species/other species, including exotic species and forest plants), and their interaction. We included plant family and genus as nested random effects to account for phylogenetic effects. We predicted that there would be a positive relationship between abundance in the two plot types if seed dispersal limited grassland species recovery in forest plots. We conducted the same analysis for each dispersal type separately and compared the native grassland species richness of each dispersal type between the two plot types using a GLM (Poisson distribution and log-link function).

We also analyzed the models in which the richness of native grassland species in each dispersal type was the response variable, whereas land-use history type and all environmental and vegetation factors were included as explanatory variables. We used R v3.3.2 and *vegan* package v2.4.1 (R core Team, 2016) for all analyses.

## Results

### Difference in vegetation and environmental factors between land use history types

Forest plots had a significantly higher MI than pasture plots (Fig. S3, Table S3, pasture plot, mean ± S.E. = 1.81 ± 0.03; forest plots, 1.91 ± 0.02), indicating that forest plots were more intensively graded than pasture plots. Slope angle did not have a significant impact on MI in either plot type (Table S3). The duration of direct sunlight was significantly shorter on the forest plots than on the pasture plots (Fig. 2; Table S3), whereas other environmental factors did not differ significantly between plot types (Fig. 2; Table S3). Furthermore, we found no significant relationships between MI and slope angle or any vegetation and environmental factors (Fig. 2; Table S3).

**Fig. 2.**
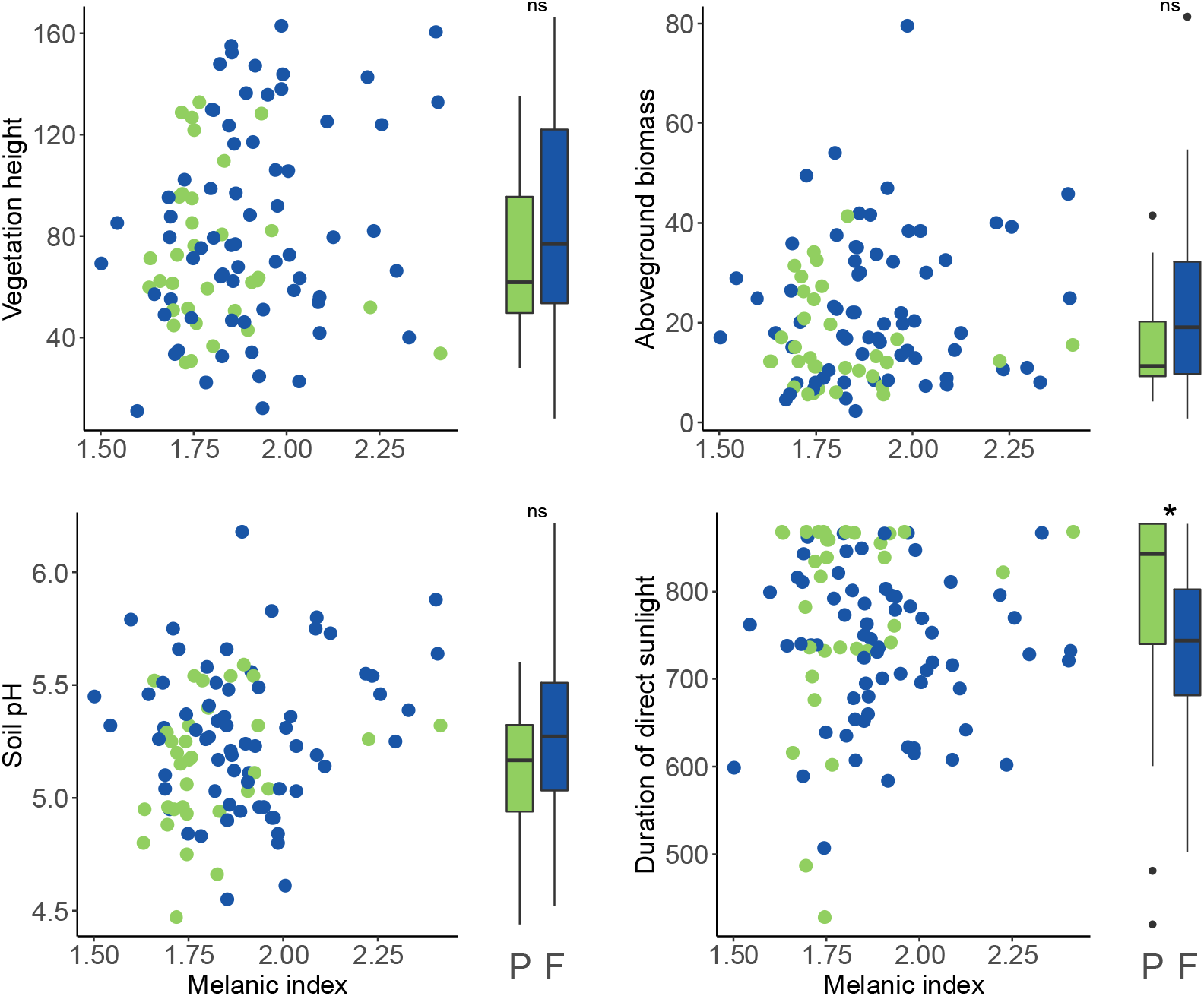
Relationships between environmental factors (vegetation height, vegetation biomass, soil pH, and duration of direct sunlight) and land use history (pasture/forest) and the intensity of machine grading (the melanic index). Green and blue circles are pasture (P) and forest (F) plots, respectively. Boxplots indicate environmental factors for P and F plots; central bars indicate the median, and asterisks indicate significant differences (*, *p* < 0.05; ns, *p* > 0.05).

### Species richness

We identified 279 plant species in the 108 studied slope plots, including 167 native grassland species, 23 exotic species, and 89 native forest species (Table S4). We detected eight endangered grassland species in the national Red List: six were vulnerable (VU), one was endangered (EN), and one was near-threatened (NT).

Total plant species (median = 36), native grassland plant species richness (27 species), and RLI (1.773) were significantly higher in pasture plots than in forest plots (median = 31 and 20 species and 1.036, respectively). Furthermore, these values declined significantly along with MI in both plot types (Fig. 3, Table S3). The occurrence of endangered grassland species was significantly higher in pasture plots than in forest plots and significantly decreased with MI in both plot types (Fig. S4, Table S3). Exotic grassland species richness was not significantly correlated with land-use history but significantly increased with MI (Fig. 3, Table S3).

**Fig. 3.**
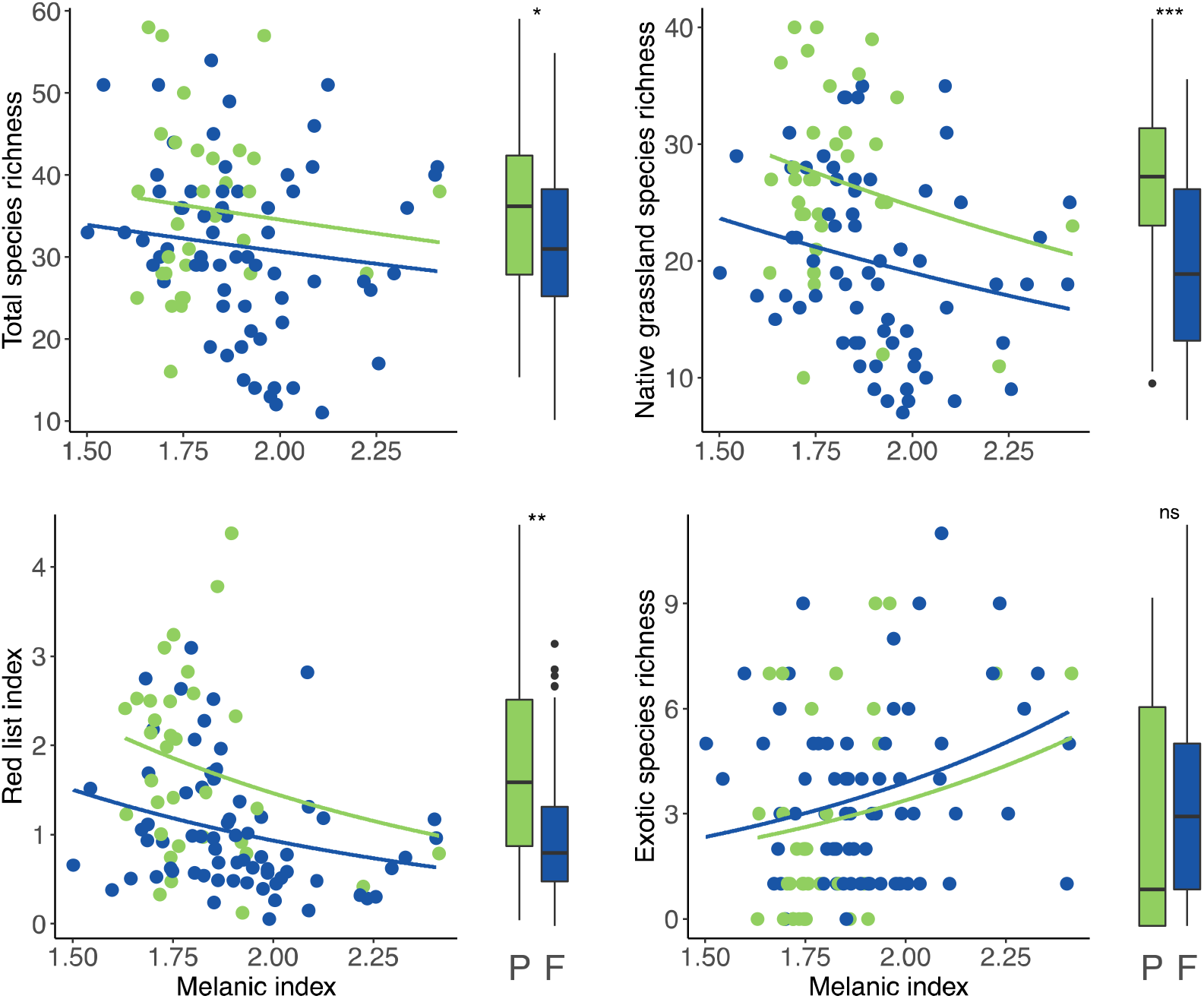
Relationships between diversity indices (total species richness; native grassland species richness; Red List Index; exotic species richness) and land use history (pasture/forest) and the intensity of machine grading (the melanic index). Green and blue circles are pasture (P) and forest (F) plots, respectively. Lines represent significant regressions estimated from GLMs. Boxplots indicate diversity indices for P and F plots; central bars indicate the median, and asterisks indicate significant differences (***, *p* < 0.001; **, *p* < 0.01; *, *p* < 0.05; ns,*p* > 0.05).

The pasture plots had significantly lower total and native grassland species richness than the reference plots at SMRC, but RLI did not differ significantly between the plot types (Table S2). When considering the forest plots, the total species richness, native grassland species richness, and RLI were significantly lower than those observed in the reference plots (Table S2).

The total species, native grassland species richness, and RLI significantly decreased with increasing aboveground biomass, whereas vegetation height did not influence any diversity indices (Table S3). Soil pH exhibited significant positive effects on all diversity indices, except exotic species richness (Table S3). Total and endangered species richness and RLI significantly increased with increasing duration of direct sunlight, whereas exotic species decreased with the variable (Table S3). Slope angle had no significant effect on any of the diversity indices (Table S3).

### Species composition

Two-dimensional NMDS ordination indicated that species composition differed substantially between pasture and forest plots (Fig. 4). Differences in centroids could not be compared because a preliminary test (using the *betadisper* function for homogeneity of multivariate dispersions) indicated a difference in the variance in species composition between the two plot types, with forest plots exhibiting a significantly higher variance than pasture plots (permutation test; *P* < 0.01; number of permutations = 999). This trend was conserved even if we considered the Euclidian distance between plots: pasture plots had a more similar species composition than forest plots (Fig. S5).

**Fig. 4.**
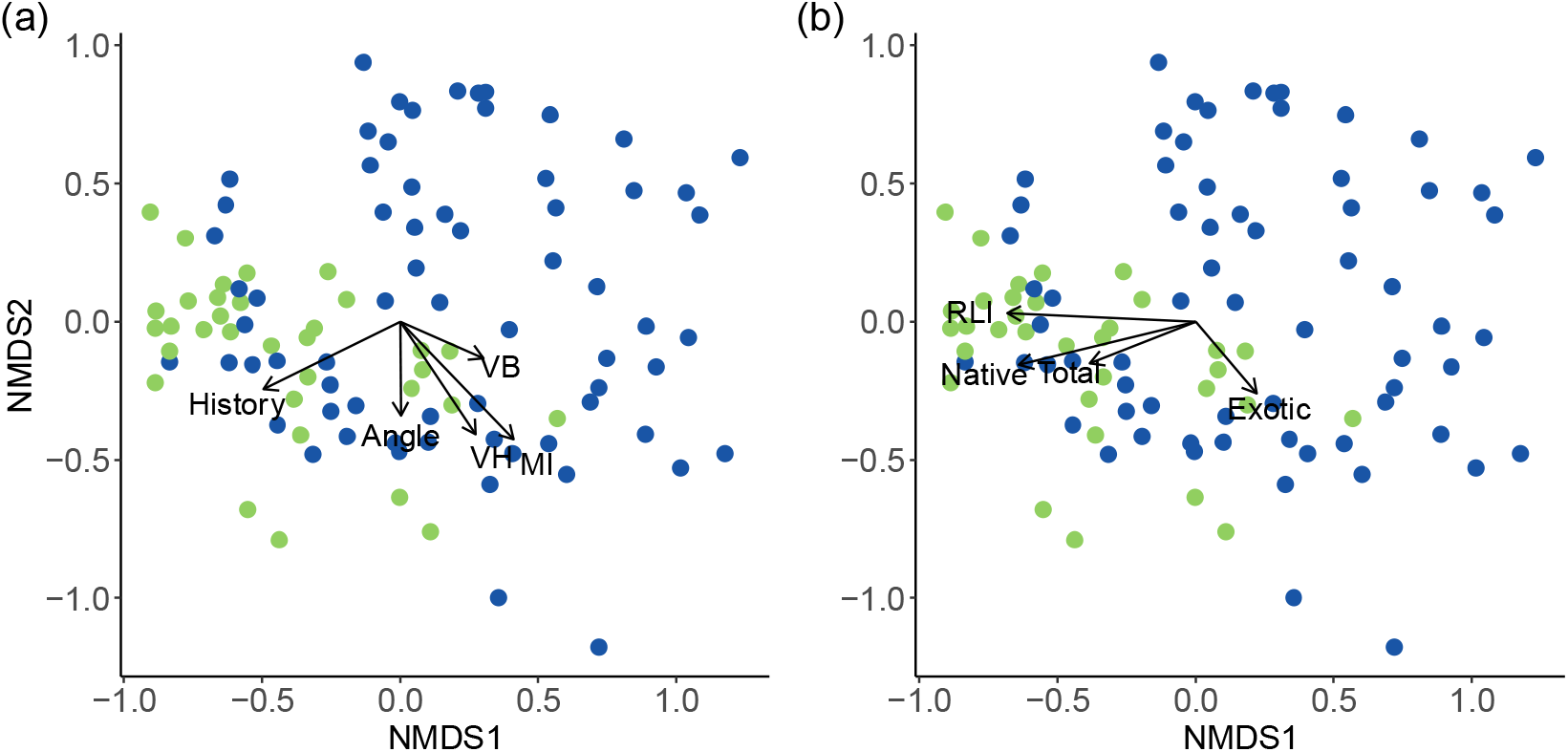
Nonmetric multidimensional scaling ordination (NMDS) plots, with significant vectors for (a) six vegetation and environmental variables (land use history (History); the Melanic Index (MI); slope angle (Angle); vegetation height (VH); aboveground biomass (AB)), and for (b) richness metrics (total (Total), native grassland (Native), and exotic (Exotic) species richness and red list index (RLI)). Arrows represent significant vectors and lengths indicate effect strength. Closed green circles and blue triangles indicate pasture and forest plots, respectively.

Vectors for land use history, MI, vegetation height, aboveground biomass, slope angle, RLI, total richness, and richness of native grassland and exotic species richness provided a significant explanation of the among-plot differences in species composition (Fig. 4). The vectors of land use history, native grassland species richness, and RLI were pointed in a similar direction, as were those of MI, vegetation height, and exotic species richness (Fig. 4).

### Species abundance and species frequency of each dispersal type

The abundance of study species in the forest plots significantly increased with their abundance in the pasture plots when considering the relationship between abundance in pasture plots and that in forest plots for each species (Fig. 5, Table S3). There was a significant interaction between species abundance in pasture plots and species group, and their abundance in forest plots. The abundance of most native grassland species was lower in forest plots than in pasture plots, and this pattern was consistent with those of barochorous and anemochorous species (Fig. S6). The abundance of other species was nearly identical between the two plot types (Fig. 5).

**Fig. 5.**
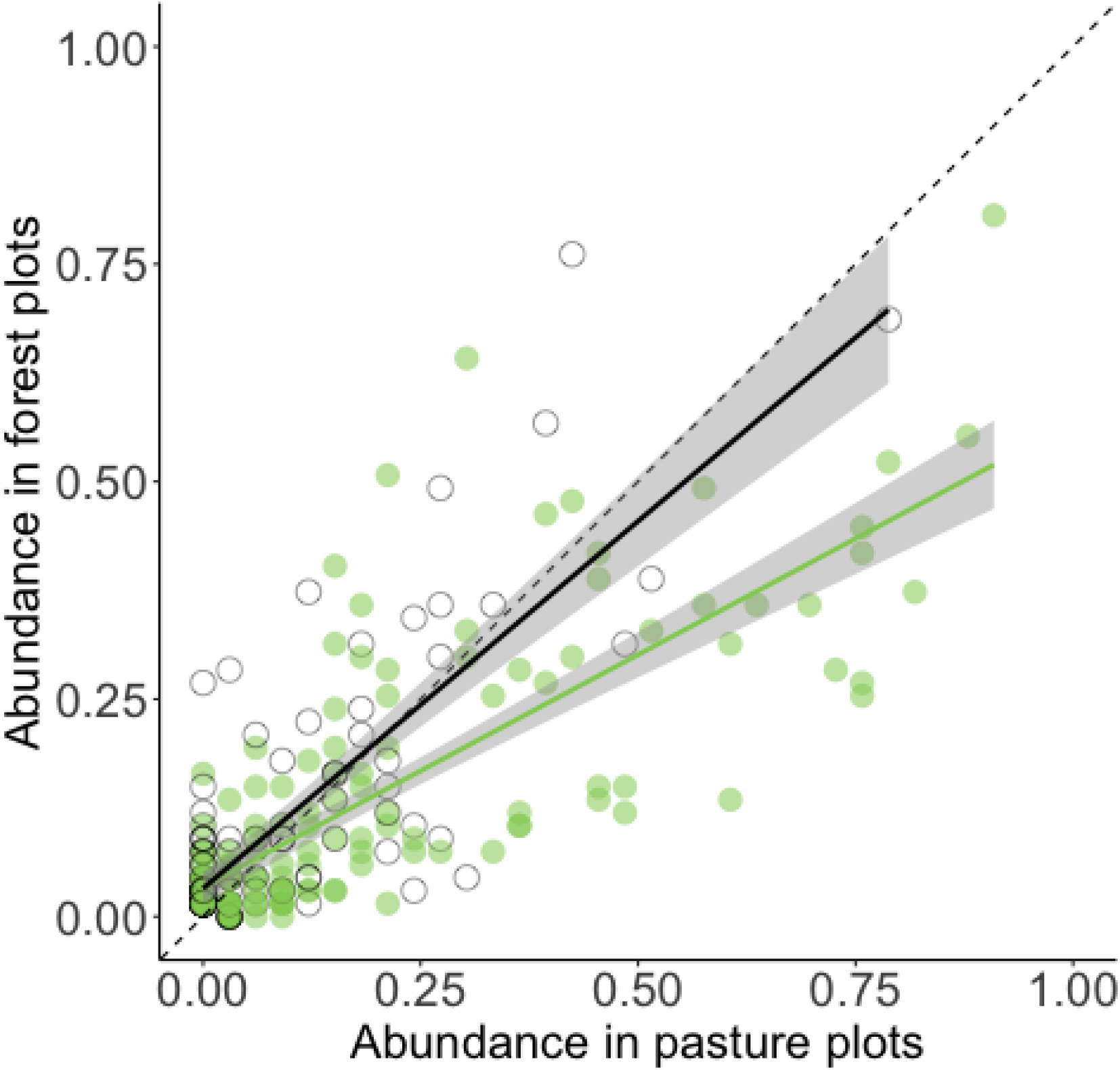
The relationship between abundance in pasture plots and that in forest plots for each species. Green circles are native grassland species, whereas black circles are other species, including forest and exotic species. The dotted line represents “y = x”, and black and green lines, and gray areas represent significant regressions and the 95 % confidential intervals estimated from the GLMM (Table S3).

The richness of barochorous and anemochorous species was significantly lower in forest plots than in pasture plots (Fig. 6). Species richness in the other dispersal types (ballochorous, hydrochory, myrmecochory, epizoochorous, and endozoochory) did not differ between plot types (Fig. 6).

**Fig. 6.**
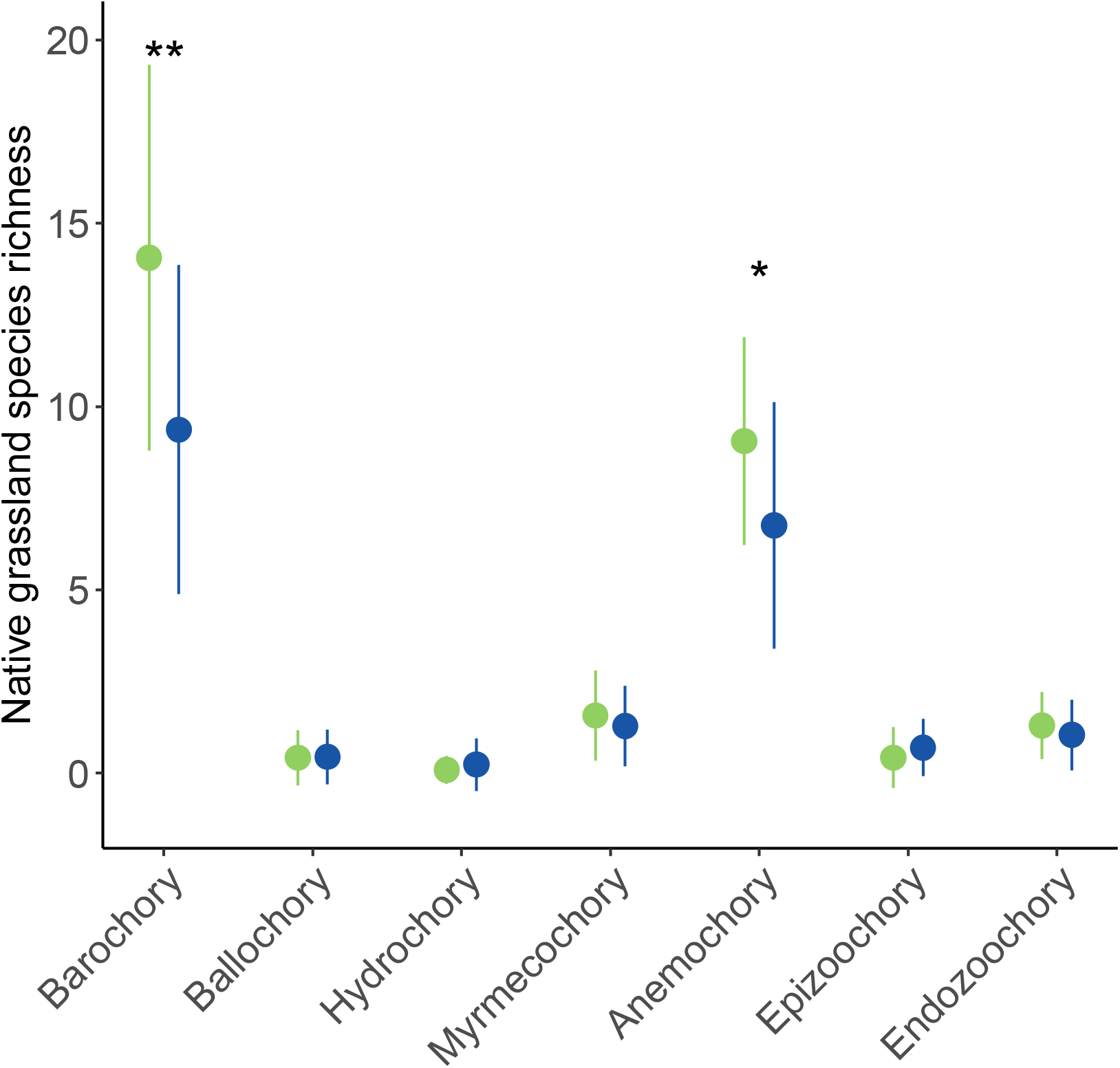
Differences in the prevalence of seven seed dispersal types between pasture (green) and forest (blue) plots for native grassland species: mean native species richness ± SD by dispersal type. Asterisks indicate significant differences between plot types (**, *p* < 0.01; *, *p* < 0.05).

## Discussion

We found that both types of ski slopes have good potential to support numerous native grassland plant species, although the native grassland species diversity on slopes was lower than that in the reference meadow. We confirmed previous findings that the potential was lowered by past forestation (Inoue et al. 2021). Two environmental factors, MI and duration of direct sunlight, were influenced by past short-term forestation, such that forest plots were significantly more graded and received less sunlight than pasture plots. Soil grading and more shaded conditions could result in negative legacy effects on the current grassland richness in forest plots. We also found a significant correlation between the abundance of each species in pasture and forest plots, and a significantly lower richness of species with lower dispersal ability in forest plots. These findings support the environmental changes and dispersal limitation hypotheses discussed in more detail below.

### Difference in vegetation and environmental factors between land use history types

We found that forest plots were more intensively graded than pasture plots, using MI as a surrogate for the intensity of machine grading on slopes. This result is plausible because machine grading is usually adopted to remove stumps and roots of trees and shrubs from ski slopes. Intense grading resulted in decreased diversity of native grassland species in both plot types, as evidenced by declines in all diversity indices, aside from exotic species richness, which conversely increased with MI. Machine grading removes the soil seedbank as well as aboveground plant material, hindering the rapid re-establishment of grassland species (Urbanska & Fattorini, 2000). Moreover, grading disturbance can lead to suitable habitats for introduced exotic grasses such as *Lolium arundinaceum, Dactylis glomerata*, and *Phleum pratense* which were more often found in forest slopes than in pasture slopes. Thus, removal of diaspore banks might have long-lasting (i.e., more than 80 years in some cases) negative legacy effects on native grassland plant diversity in the study forest plots (Wipf et al., 2005; Roux-Fouillet et al., 2011).

Compared to pasture plots, forest plots received a shorter duration of direct sunlight, which caused a lower incidence of endangered species and a higher diversity of exotic species (c.f., Yaida et al., 2019). Forest slopes are usually narrower than pasture slopes, likely because of the cost of deforestation during slope construction (Fig. 1). Poor light environments due to shading by surrounding forests might partly affect the negative legacy effects on forest slopes. Thus, our findings support the hypothesis of environmental change such that increased grading and shading have caused lower grassland species in forest slopes than in pasture slopes.

### Dispersal limitation explaining low grassland species diversity in forest plots

The recovery of grassland communities, including the re-establishment of endangered species, might be limited even after more than 50 years post ski resort construction on forest slopes. We previously reported that forest understory communities adjacent to ski slopes supported only a few grassland species at very low abundance (Inoue et al., 2021). Grassland species have been shown to have poor seed banking capacity in soils in Europe and Japan (Russi, Cocks, & Roberts, 1992; Koyanagi et al., 2011). We also confirmed that few species made seed banks on the study slopes and nearby forests (Fig. S7). Moreover, forest slopes are often more graded, as we have shown. Thus, most grassland species were once eliminated on forest slopes, and their re-colonization and/or re-establishment might depend on seed dispersal from neighboring grasslands, which would also explain the large differences in species composition between the plot types.

Indeed, we found significantly lower richness of native species in the two major dispersal types (barochory and anemochory) in forest plots than in pasture plots. Conversely, the richness of animal-dispersed species in forest plots was nearly equal to that in pasture plots for myrmecochorous, endozoochorous, and epizoochorous species. Barochorous species usually have limited dispersal ability compared to animal-dispersed species (Thomson, Moles, Auld, & Kingsford, 2011). Thus, dispersal limitation is hypothesized to be the cause of lower grassland species richness observed on forest slopes. The abundance of native grassland species in forest plots was positively correlated with their abundance on pasture slopes, which are likely to serve as a seed source, supporting our hypothesis that more abundant species will have a higher colonizing success.

The dispersal limitation hypothesis can also explain the significant, large inter-plot vegetation dissimilarity (beta diversity) on forest slopes compared to pasture plots (Fig. 4 and permutation test; *P* < 0.01). If grassland recovery on forest slopes is limited by seed dispersal rather than by environmental filtering, we would expect species composition to be strongly shaped by stochastic re-colonization, resulting in high beta diversity (Freestone & Inouye, 2006).

However, one result is more difficult to interpret. The richness of anemochorous species, which often have good long-distance dispersal capacities (Thomson, 2011), is lower on forest slopes than on pasture slopes. This might be attributable to the fact that there were many anemochorous species that were rare in the pasture plots and those with relatively low dispersal ability (Fig. S6).

### Implications of the study for grassland plant conservation on ski slopes

This study, along with previous studies, demonstrates that ski resort development can maintain semi-natural grassland habitats and sustain biodiversity below the treeline. Echoing results from other studies worldwide, our findings indicate there can be positive conservation outcomes from ski resort developments for threatened semi-natural grassland species (Lanvers, Sieg, & Fartmann, 2012; Erfanian et al., 2019; Yaida et al., 2019). However, we also found that the potential for ski slopes to provide habitat for grassland species may be depressed by past short-term forest use, concomitant intensive grading, and poor light environments during and after slope construction. Grassland species with zoochorous dispersal and those with higher abundance on pasture slopes more frequently recolonized forest slopes, suggesting that dispersal limitations affect the re-establishment of grassland communities on such ski slopes. To mitigate this limitation, sowing or planting native grassland species present in neighboring pasture slopes to restore species-rich grassland conditions on forest slopes rather than introducing exotic grasses is a potential solution.

Unfortunately, numerous ski resorts have been abandoned in the last few decades in Japan for socioeconomic reasons (c.f., Nakamura, 1999), and over-machine grading and ornamental flower planting (for summer sightseers) are often used in active ski resorts (Yaida et al., 2019). Conservation of biodiversity-rich pasture slopes and restoration of forest slopes as habitats for threatened native grassland species can be promoted through the maintenance of annual mowing practices, avoidance of intensive machine grading, and wider ski courses. If grading is inevitable, it is advisable to stockpile topsoil, which contains propagules of native grassland species, and replace it in situ following grading. Our findings could be useful to resort companies, national and local governments, citizens, conservationists, and researchers, who may assume that ski development always negatively impacts mountain ecosystems and their ecosystem services.

## Supporting information

Supporting imformation

## Abbreviations

SMRC: Sugadaira Montain Research Center
EN: endangered
VU: vulnerable
NT: near threatened
RLI: red list index
MI: Melanic Index
JDI: Jaccard dissimilarity index

## Acknowledgements

We thank T. Inoue, Y. Sakata, N. Nakahama, S. Matsuhisa, K. Aoki, K. Murakami, K. Takashima, T. Seki for their field assistance, A. Nikkeshi and M. K. Hiraiwa for their suggestions on statistical analyses, T. Inoue for his help in the GIS analysis, and Y. Iimura for helping the humic acid analysis. We are grateful to land owners of our study sites, which are “general foundation Nirei kai”, “Sugadaira Bokujo”, ”Hatsune-kan ”, “Imai-kan”, “Jozan-kan”, “Shinshu Yunomaru Snow Resort”, and “Kazawa Snow Area”; and the management companies, which are “Sugadaira Pine beak ski“, “Sugadaira Ski House Co., Ltd.”, “Oku Davos Snow Park”, “HARE Sugadaira-Kogen Snow Resort”, “Yunomaru-kanko-kaihatsu Co., Ltd.”, and “Gorin-kanko Co., Ltd.”; for their allowing our field surveys. This work was supported by JSPS KAKENHI Grant Nos. 16H02993, 17K07557, and 19H03303.

## Authors’ contributions

AU and TK conceived the idea of this work; AU, YAY, KU and TK designed the study; All authors collected the data; YAY and AU analyzed the data and wrote the first draft. All authors contributed to the drafts and gave final approval for publication. YAY, TK, and AU contributed mainly and equally to the work.

## References

Burt, J. W., & Rice, K. J. 2009. Not all ski slopes are created equal: Disturbance intensity affects ecosystem properties. Ecological Applications, 19(8), 2242–2253.

Butchart, S. H. M., Walpole, M., Collen, B., Strien, A. van, Scharlemann, J. P. W., Almond, R. E. A.,… Watson, R. 2010. Global Biodiversity: Indicators of Recent Declines. Science, 1187512.

Dormann, C. F., McPherson, J. M., Araújo, M. B., Bivand, R., Bolliger, J., Carl, G.,… Wilson, R. 2007. Methods to account for spatial autocorrelation in the analysis of species distributional data: a review. Ecography, 30(5), 609–628.

Erfanian, M. B., Ejtehadi, H., Vaezi, J., Moazzeni, H., Memariani, F., & Firouz-Jahantigh, M. 2019. Plant community responses to environmentally friendly piste management in northeast Iran. Ecology and Evolution, 9(14), 8193–8200.

Foley, J. A., DeFries, R., Asner, G. P., Barford, C., Bonan, G., Carpenter, S. R.,… Snyder, P. K. 2005. Global Consequences of Land Use. Science, 309(5734), 570–574.

Freestone, A. L., & Inouye, B. D. 2006. Dispersal Limitation and Environmental Heterogeneity Shape Scale-Dependent Diversity Patterns in Plant Communities. Ecology, 87(10), 2425–2432.

Gerstner, K., Dormann, C. F., Stein, A., Manceur, A. M., & Seppelt, R. 2014. Effects of land use on plant diversity – A global meta-analysis. Journal of Applied Ecology, 51(6), 1690–1700.

Gunma prefecture, 2018. Red Data Book Gunma edition - Plants. Gunma Prefecture, Gunma, Japan. (in Japanese)

Honna, T., Yamamoto, S., & Matsui, K. 1988. A Simple Procedure to Determine Melanic Index that is Useful for Differentiating Melanic from Fulvic Andisols. Pedologist, 32(1), 69–78.

Inoue, T., Yaida, Y. A., Uehara, Y., Katsuhara, K. R., Kawai, J., Takashima, K., Ushimaru, A., & Kenta, T. 2021. The effects of temporal continuities of grasslands on the diversity and species composition of plants. Ecological Research. 36(1), 24–31.

Kawano, T., Sasaki, N., Hayashi, T., & Takahara, H. 2012. Grassland and fire history since the late-glacial in northern part of Aso Caldera, central Kyusyu, Japan, inferred from phytolith and charcoal records. Quaternary International, 254, 18–27.

Koyanagi, T., Kusumoto, Y., Yamamoto, S., Okubo, S., Kitagawa, Y., & Takeuchi, K. 2011. Potential for restoring grassland plant species on an abandoned forested Miscanthus grassland using the soil seed bank as a seed source. Japanese Journal of Conservation Ecology, 16(1), 85–97. (in Japanese)

Koyanagi, T. F., & Furukawa, T. 2013. Nation-wide agrarian depopulation threatens semi-natural grassland species in Japan: Sub-national application of the Red List Index. Biological Conservation, 167, 1–8.

Lampinen, J., K. Ruokolainen, and A.-P. Huhta. 2015. Urban Power Line Corridors as Novel Habitats for Grassland and Alien Plant Species in South-Western Finland. PLoS ONE, 10.

Lanvers, J., Sieg, B., & Fartmann, T. 2012. Impacts of cross-country ski trails on bog and fen vegetation. Tuexenia, 32, 87–103.

Ministry of the Environment of Japan. 2019. The 4^th^ version of the Japanese red list on 9 taxonomic groups. Ministry of the Environment Government of Japan, Tokyo. (in Japanese)

Miyabuchi, Y., Sugiyama, S., & Nagaoka, Y. 2012. Vegetation and fire history during the last 30,000 years based on phytolith and macroscopic charcoal records in the eastern and western areas of Aso Volcano, Japan. Quaternary International, 254, 28–35.

Nagata, Y. K., & Ushimaru, A. 2016. Traditional burning and mowing practices support high grassland plant diversity by providing intermediate levels of vegetation height and soil pH. Applied Vegetation Science, 19(4), 567–577.

Nagano Prefecture, 2014. Red Data Book Nagano edition-Plants. Nagano Prefecture, Nagano Japan. (in Japanese)

Nakahama, N., Uchida, K., Ushimaru, A., & Isagi, Y. 2018. Historical changes in grassland area determined the demography of semi-natural grassland butterflies in Japan. Heredity, 121(2), 155.

Nakamura, T. 1999. A review of scientific studies on ski areas in Japan. Japanese Journal of Ecology, 49, 261–264. (in Japanese)

Newbold, T., Hudson, L. N., Hill, S. L. L., Contu, S., Lysenko, I., Senior, R. A.,… Purvis, A. 2015. Global effects of land use on local terrestrial biodiversity. Nature, 520(7545), 45–50.

Ogura, J. 2006. The transition of grassland area in Japan. Journal of Kyoto Seika University 30, 160–172. (in Japanese)

Ohara, R. G., & Ushimaru, A. 2015. Plant beta-diversity is enhanced around grassland–forest edges within a traditional agricultural landscape. Applied Vegetation Science, 18(3), 493–502.

Öster, M., Ask, K., Cousins, S.A.O., & Eriksson, O. 2009. Dispersal and establishment limitation reduces the potential for successful restoration of semi-natural grassland communities on former arable fields. Journal of Applied Ecology, 46, 1266–1274.

R Core Team. 2016. R: A language and environment for statistical computing. R Foundation for Statistical Computing, Vienna, Austria. URL: https://www.R-project.org/.

Rolando, A., Caprio, E., Rinaldi, E., & Ellena, I. 2007. The impact of high-altitude ski-runs on alpine grassland bird communities. Journal of Applied Ecology, 44(1), 210–219.

Roux-Fouillet, P., Wipf, S., & Rixen, C. 2011. Long-term impacts of ski piste management on alpine vegetation and soils. Journal of Applied Ecology, 48(4), 906–915.

Russi, L., Cocks, P. S., & Roberts, E. H. 1992. Seed Bank Dynamics in a Mediterranean Grassland. Journal of Applied Ecology, 29(3), 763–771.

Sala, O. E., Chapin, F. S., Iii, Armesto, J. J., Berlow, E., Bloomfield, J.,… Wall, D. H. 2000. Global Biodiversity Scenarios for the Year 2100. Science, 287(5459), 1770–1774.

Satake, Y., Ohwi, J., Kitamura, S., Watari, S. & Tominari, T. 1981. Wild Flower of Japan: Herbaceous Plants. Vols. I–III. Heibon-sha, Tokyo, JP. (in Japanese)

Takahashi, T. & Shoji, S. 2002. Distribution and classification of volcanic ash soils, Global Environmental Research, 6(2), 83–97.

Thomson, F. J., Moles, A. T., Auld, T. D., & Kingsford, R. T. 2011. Seed dispersal distance is more strongly correlated with plant height than with seed mass. Journal of Ecology, 99(6), 1299–1307.

To□ro□k, P. & Dengler, J. 2018. Palaearctic grasslands in transition: overarching patterns and future prospects. In: Squires, V.R., Dengler, J., Feng, H. & Hua, L. (eds.) Glasslands of the world: diversity, management and conservation, pp. 15–26. CRC Press, Boca Raton, US.

Tsuyuzaki, S. 1990. Species composition and soil erosion on a ski area in Hokkaido, northern Japan. Environmental Management, 14(2), 203–207.

Tsuyuzaki, S. 1994. Environmental Deterioration Resulting from Ski-resort Construction in Japan. Environmental Conservation, 21(2), 121–125.

Tsuyuzaki, S. 1995. Ski slope vegetation in central Honshu, Japan. Environmental Management, 19(5), 773.

Tsuyuzaki, S. 2002. Vegetation development patterns on skislopes in lowland Hokkaido, northern Japan. Biological Conservation, 108(2), 239–246.

Uchida, K., & Ushimaru, A. 2015. Land abandonment and intensification diminish spatial and temporal ß-diversity of grassland plants and herbivorous insects within paddy terraces. Journal of Applied Ecology, 52(4), 1033–1043.

Uchida, K., Takahashi, S., Shinohara, T., & Ushimaru, A. 2016. Threatened herbivorous insects maintained by long-term traditional management practices in semi-natural grasslands. Agriculture, Ecosystems & Environment, 221, 156–162.

Ushimaru, A., Uchida, K. & Suka, T. 2018. Grassland biodiversity in Japan: threats, management and conservation. In: Squires, V.R., Dengler, J., Feng, H. & Hua, L. (Eds.), Grasslands of the world: diversity, management, and conservation (pp. 197–218). CRC Press, Boca Raton, US.

Ushimaru A, Uchida K, Ikegami M and Suka T. in press. Grasslands and shrublands in Japan. In Goldstein MI & DellaSala L (Eds.), The Encyclopedia of the World’s Biomes. Elsevier, Oxford.

Urbanska, K. M., & Fattorini, M. 2000. Seed Rain in High-Altitude Restoration Plots in Switzerland. Restoration Ecology, 8(1), 74–79.

Yaida, Y. A., Nagai, T., Oguro, K., Katsuhara, K. R., Uchida, K., Kenta, T., & Ushimaru, A. 2019. Ski runs as an alternative habitat for threatened grassland plant species in Japan. Palaearctic Grasslands, 42, 16–22.

Yamamoto, S., Honna, T., & Utsumi, K. 2000. Easy estimation of humic acid types by using melanic index. Journal of the science of soil and manure, Japan, 71(1), 82–85. (in Japanese)

Yasuda, M., and F. Koike. 2006. Do golf courses provide a refuge for flora and fauna in Japanese urban landscapes? Landscape and Urban Planning, 75:58–68.

Watson, A. 1985. Soil erosion and vegetation damage near ski lifts at Cairn Gorm, Scotland. Biological Conservation, 33(4), 363–381.

Wipf, S., Rixen, C., Fischer, M., Schmid, B., & Stoeckli, V. 2005. Effects of ski piste preparation on alpine vegetation. Journal of Applied Ecology, 42(2), 306–316.

