## Supporting imformation for "Negative legacy effects of past forest use on plant diversity in semi-natural grasslands on ski slopes"

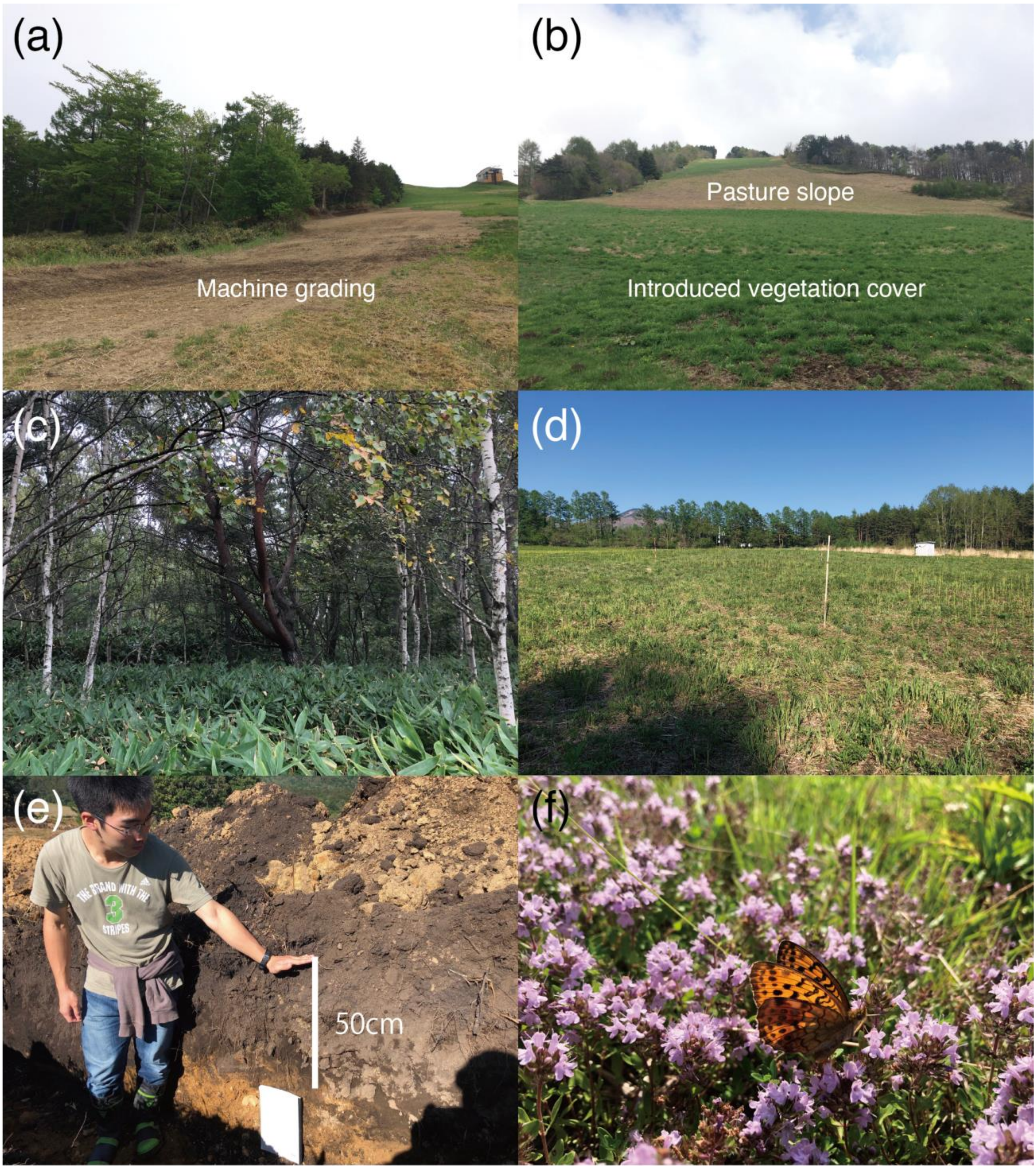

Fig. S1. Photographs of (a) a machine graded forest slope, (b) a pasture slope with introduced vegetation cover after grading (lower half), (c) *Sasa*-dominated forest floor in a secondary forest established on an old pasture, (d) a reference grassland at SMRC, (e) *Kurobokudo* topsoil on a pasture slope, and (f) *Brenthis daphne* visiting flowers of the native grassland species *Thymus quinquecostatus*, in a pasture slope community.

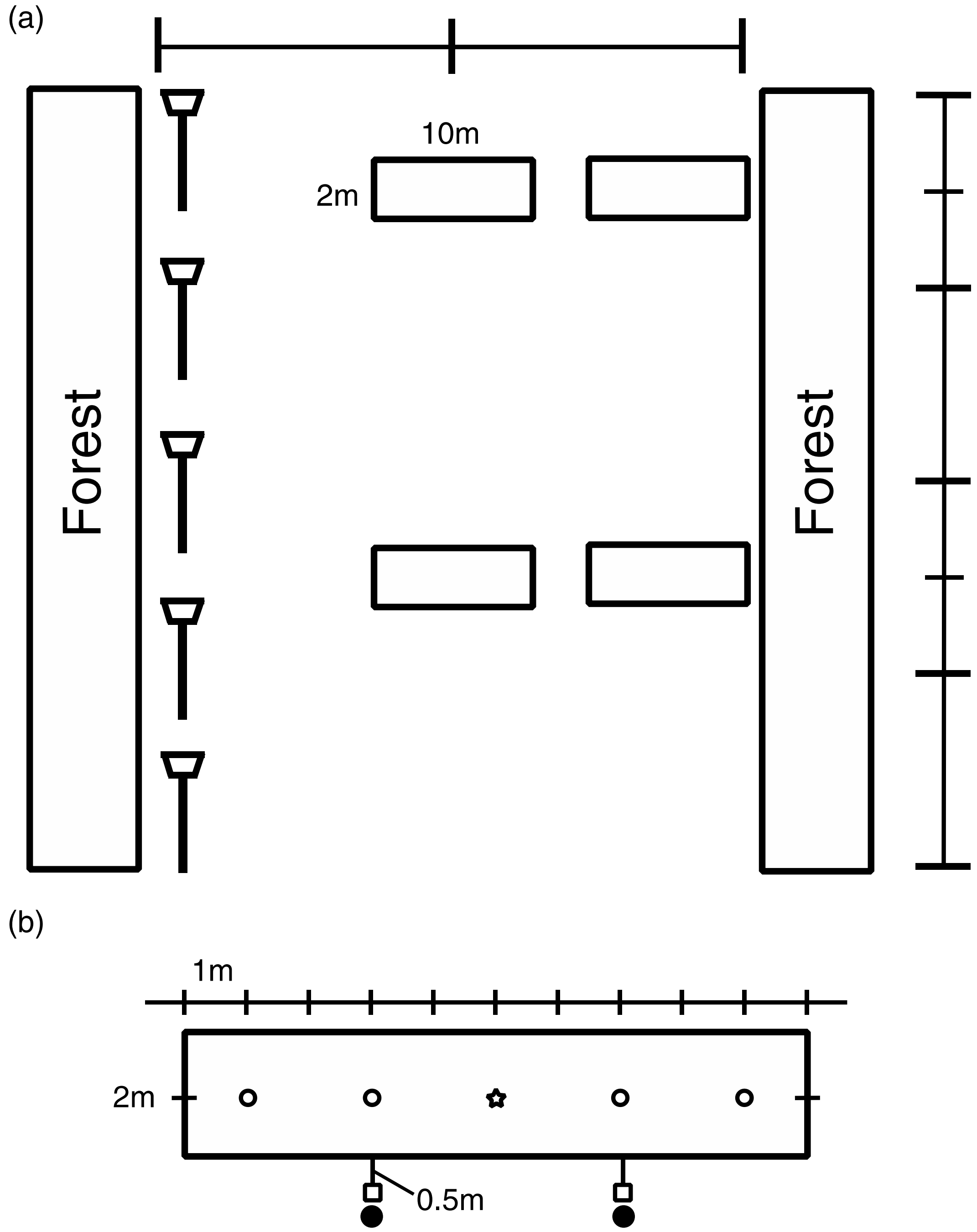

Fig. S2. Sample plot layout (a) and sampling points for environmental factors within plots (b). We first divided each slope into four segments from top to the bottom, and then established four 2 × 10 $m^2$ plots at the center and edge in the first and third segments of each slope, opposite the lift (a). In each plot, vegetation height was assessed at five points (open circles and star), and a hemispherical photograph was taken at 1 m height from the ground at the center (star). Aboveground biomass was examined in two 0.25 × 0.25 $m^2$ subplots adjacent to the plot (squares), while soils were sampled adjacent to subplots (closed circles) (b).

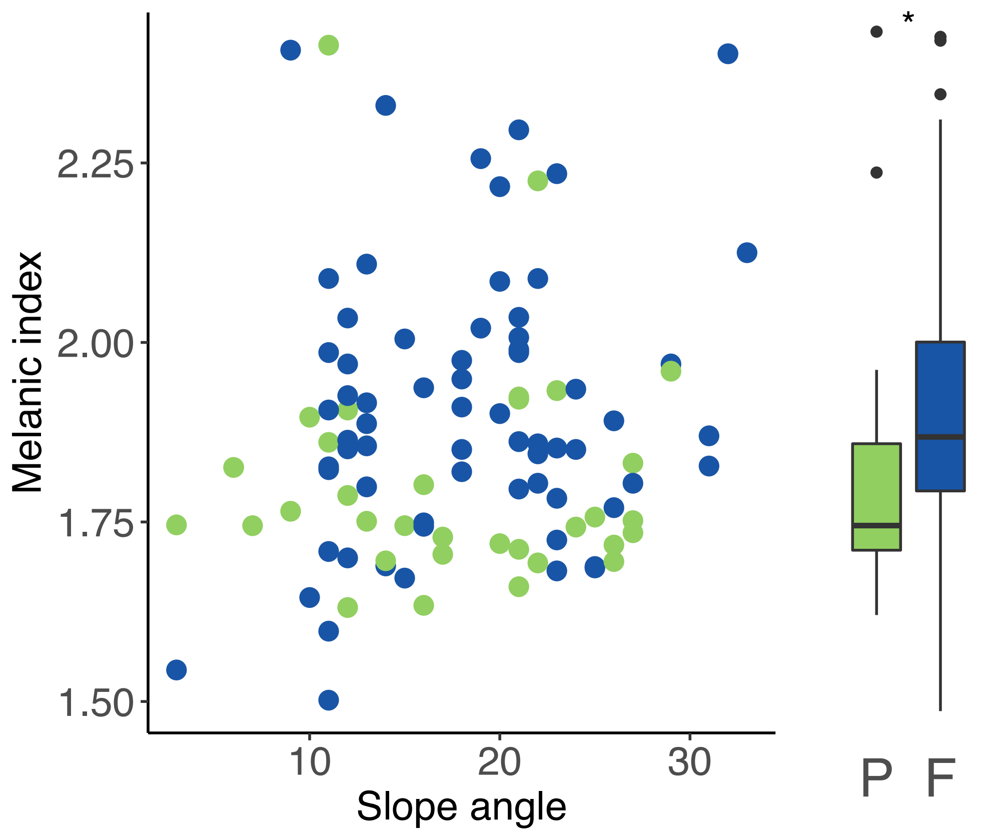

Fig. S3. Relationships between intensity of machine grading (the melanic index) and land use history (pasture/forest) and slope angle, as determined in the GLM. Green and blue circles are pasture (P) and forest (F) plots, respectively. Estimated values are listed in Table S3.

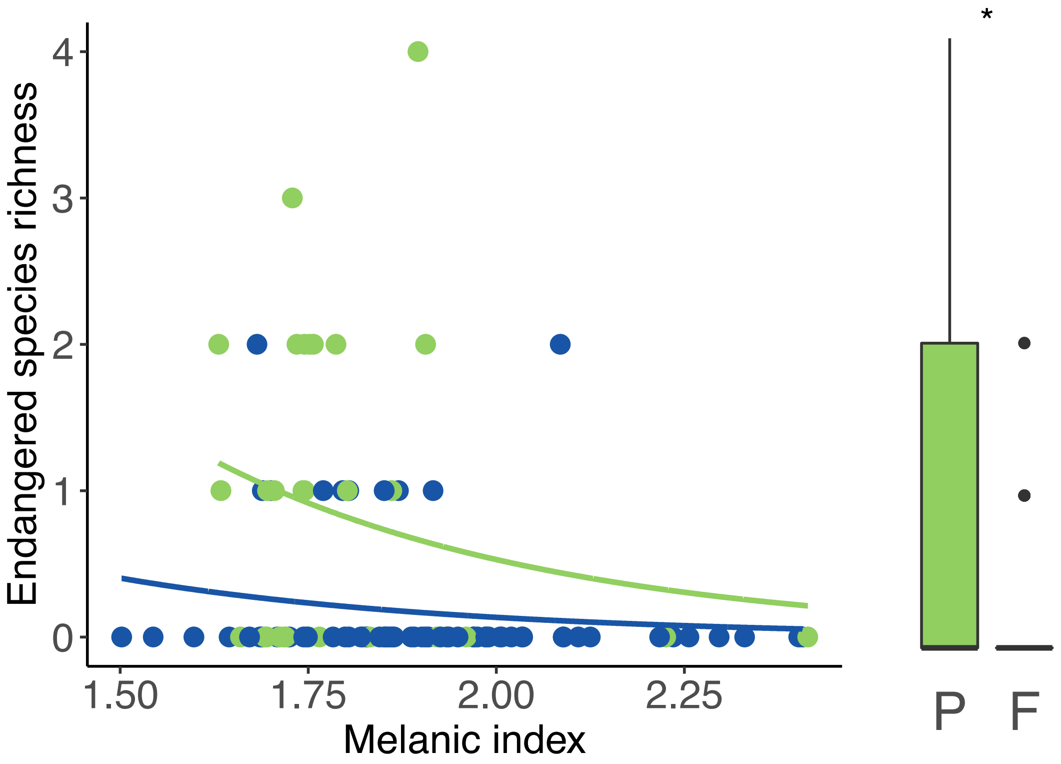

Fig. S4. Relationships between endangered species richness and land use history (pasture/forest) and the intensity of machine grading (the melanic index), as determined in the GLM. Green and blue circles are pasture (P) and forest (F) plots, respectively. Endangered species were defined as those listed on the national Red List with ranks of near-threatened or higher. Estimated values are listed in Table S3.

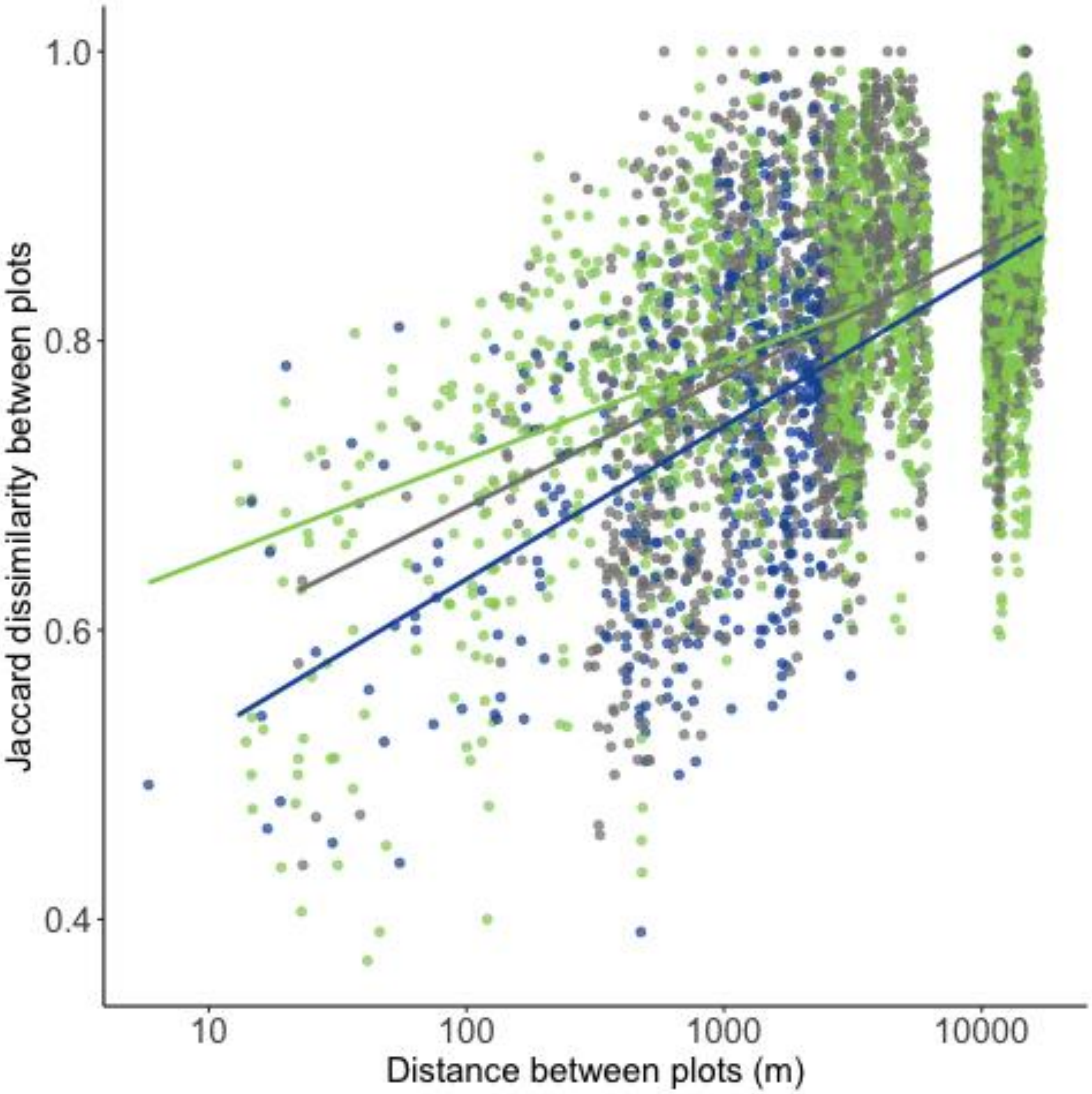

Fig. S5. The relationship between Jaccard dissimilarity and log-transformed distance (Euclidean distance; m) between plots. Plot type pairs are indicated by different colors: green = pasture-pasture, blue = forest-forest, and grey = pasture-forest. Because NMDS does not consider the effects of spatial autocorrelation on vegetation dissimilarity, we analyzed the effects of Euclidean distance (m) on vegetation differences within and between each land use history type using a GLM with a Gaussian distribution and an identity-link function. In the GLM, JDI for each plot pair was the response variable, while explanatory variables were log-transformed distance (m), Bray-curtis dissimilarity of six environmental factors, and pair type (pasture-pasture, pasture-forest, or forest-forest), and the interaction between the distance and the dissimilarity to pair type. An exact permutation test was used to assess the significance of each explanatory variable (Table S3).

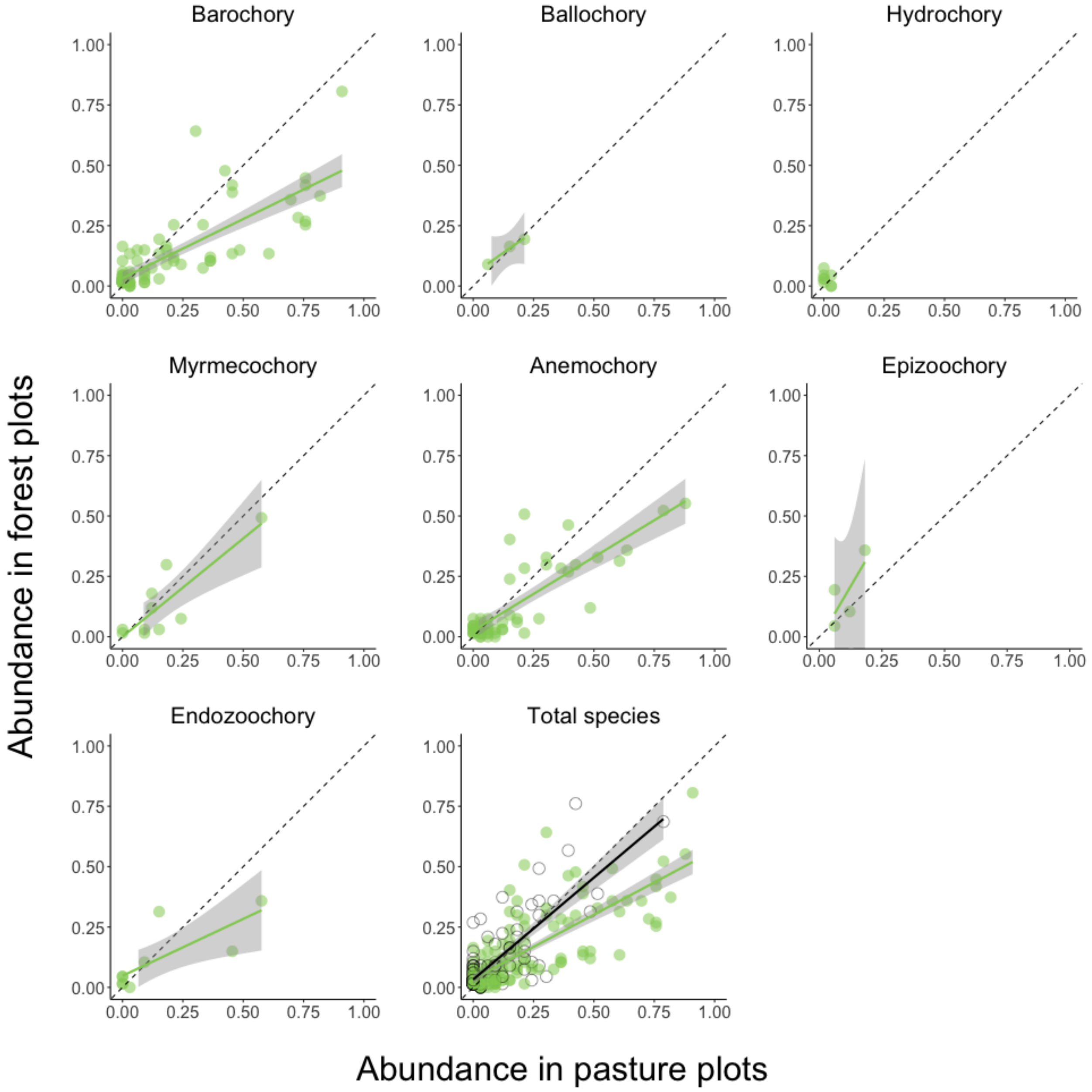

Fig. S6. Relationship between abundance in pasture plots and abundance in forest plots by dispersal type. Black circles indicate forest and exotic species, and green circles are native grassland species. The dotted line represents “y = x”, and black and green lines represent significant regressions estimated in the GLMMs. All regressions other than hydrochorous species were significant.

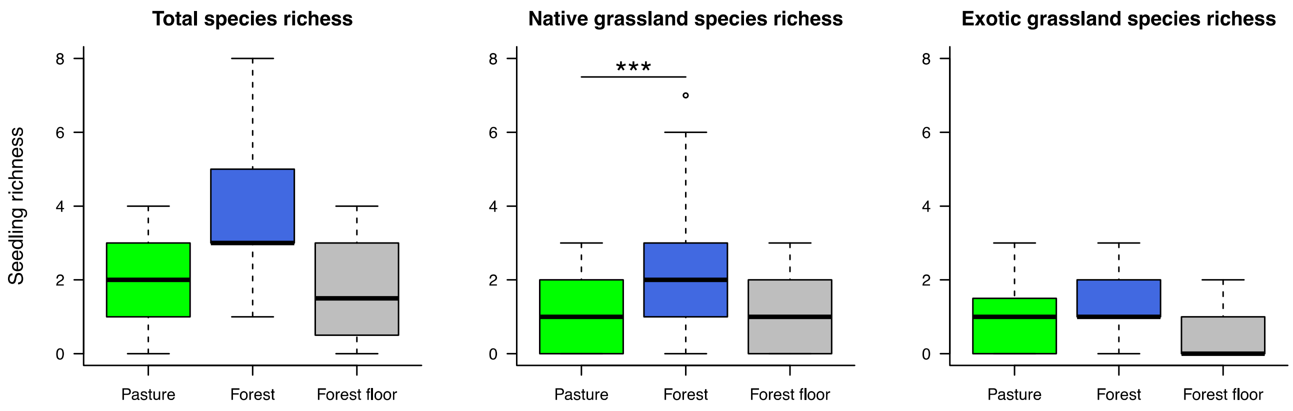

Fig. S7. Differences in richness of total, native grassland, and exotic grassland species seedlings germinated from soils of pasture and forest plots and adjacent forest floors (PS: pasture plot; FS: forest plot; AF: adjacent forest floor plot). Significant difference between slope types is indicated by asterisks (***, $p < 0.001$).

Methods: Seed bank survey was conducted on 20 ski slopes and 20 adjacent forest floors in 5 ski resorts (6, 7, 3, 2 and 2 slopes in Dabosu, Taro, Omatsu, Tsubakuro, and Minenohara ski resorts, respectably), and we set 2 plots on upper and lower of each ski slope and 1 plots in a adjacent forest for each slope: we totally set 40 slope plots and 20 forest floor plots. We collected two soil samples in each plot on May 2018, using a cylindrical soil corer with a diameter of 5-cm and a height of 5-cm (100 $cm^3$ per sample and 12,000 $cm^3$ soil in total). Living materials, stones and coarse roots were removed from each soil sample. Then, each soil sample was planted in pots filled with mixed vermiculite and red granular soil (1:1) in a greenhouse at Kobe University, Japan. We watered all planted pots every day. Germinations of seedlingls were monitored and recorded from late May to early November. We pooled data from two soil samples for each plot for the following analyses,.

Results: From the 120 soil samples, we recorded seedling germinations of 37 species including 28 native and 7 exotic grassland species and a native forest species seedlings were also recorded. Thus, we found only a limited number of germinated grassland species from soil samples. We totally observed 22, 33, and 18 species on pasture and forest slopes and adjacent forest floors, respectively. Seedling richness of native grassland species on pasture and adjacent forest floor plots were lower (median richness, 1.5 and 1.5 species, respectively; Fig. 1) than forest slope (median richness, 2.0 species; Fig. 1), although significant difference was found only between pasture and forest plots. Seedling richness of total and exotic grassland species did not differ between plot types.

Table S1. The number of study slopes (plots) in each land use history type and slope area, elevation range (minimum and maximum), and latest observation of forest in forest plots at each ski resort. Slopes within the same resort are located on/in the same mountain ridge or valley.

| | Number of slopes (plots) | | Area (min.-max.; ha) | Elevation (min.-max.; m) | Latest observation of forest in forest plots |
|---|---|---|---|---|---|
| Area name | Pasture | Forest | | | |
| Sugadaira-plateau | | | | | |
| Dabosu-yama | 5 (18) | 3 (10) | 2.09-7.70 | 1330-1520 | 1910 |
| Tarou-yama | 3 (7) | 6 (17) | 1.96-4.22 | 1270-1440 | 1975-1965 |
| Omatsu-yama | 0 | 2 (8) | 1.62-2.08 | 1380-1610 | 1947-1991 |
| Tsubakuro-yama | 0 | 1 (4) | 0.99 | 1340-1470 | 1937-1975 |
| Minenohara-plateau | 2 (8) | 1 (4) | 1.72-3.37 | 1400-1530 | 1965 |
| Yunomaru-plateau | 0 | 3 (12) | 4.14-11.06 | 1730-1900 | 1947-1991 |
| Kazawa-plateau | 0 | 3 (12) | 1.04-3.07 | 1350-1640 | 1947-1991 |
| Total | 10 (33) | 19 (67) | | | |

Table S2. Plant species richness and Red List Index (RLI) of pasture, forest, and reference meadow plots on historical pastures (SMRC, Tsukuba University). Significant differences between reference and other plot types were assessed using GLMs with a Poisson distribution for richness and a Gaussian distribution for RLI: *** $p < 0.001$; * $p < 0.05$; no annotation, $p > 0.05$.

| | No. of | Species richness (mean ± SE) | | | |
|---|---|---|---|---|---|
| Types | plots | Total | Native grassland | Exotic grassland | RLI |
| Reference meadow | 8 | 41.6 ± 1.2 | 35.8 ± 1.0 | 4.5 ± 0.9 | 1.7 ± 0.2 |
| Ski slope grassland | | | | | |
| Pasture | 33 | 35.9 ± 1.3* | 26.9 ± 1.0*** | 2.8 ± 0.4* | 1.8 ± 0.1 |
| Forest | 67 | 31.3 ± 1.7*** | 19.9 ± 1.3*** | 3.6 ± 0.4 | 1.0 ± 0.1* |

Table S3. GLM and Wald test results for six vegetation and environmental factors (vegetation height, aboveground biomass, soil pH, duration of direct sunlight, and slope angle), and total, native grassland and exotic species richness, as well as RLI and the abundance of native grassland species and other species in pasture plots. Estimated coefficients, SE, *z* or *t* values and *p* values for intercept and each explanatory variable are shown. Statistically significant *p* values are bolded.

**Generalized liner models**

| **Predictors** | **Estimate** | **SE** | ***t*** | ***P*** |
|---|---|---|---|---|
| Response: Melanic index | | | | |
| **Intercept** | 1.747 | $7.442 \times 10^{-2}$ | 23.474 | **<0.001** |
| **Land use history** | $9.787 \times 10^{-2}$ | $3.906 \times 10^{-2}$ | 2.506 | **<0.05** |
| Slope angle | $3.167 \times 10^{-3}$ | $2.971 \times 10^{-3}$ | 1.066 | 0.289 |
| The spatial autocovariate | $1.504 \times 10^{-7}$ | $2.564 \times 10^{-6}$ | 0.059 | 0.953 |
| Response: Vegetation height | | | | |
| Intercept | - 3.610 | $3.775 \times 10$ | - 0.096 | 0.924 |
| Land use history | 7.892 | 7.985 | 0.988 | 0.326 |
| Melanic index | $2.477 \times 10$ | $2.023 \times 10$ | 1.225 | 0.224 |
| Slope angle | $5.853 \times 10^{-1}$ | $5.926 \times 10^{-1}$ | 0.988 | 0.326 |
| **The spatial autocovariate** | $2.060 \times 10^{-5}$ | $8.927 \times 10^{-6}$ | 2.307 | **<0.05** |
| Response: Aboveground biomass | | | | |
| Intercept | 3.907 | $1.390 \times 10$ | 0.281 | 0.779 |
| Land use history | 5.786 | 2.930 | 1.975 | 0.051 |
| Melanic index | 2.906 | 7.373 | 0.394 | 0.694 |
| Slope angle | $2.280 \times 10^{-1}$ | $2.153 \times 10^{-1}$ | 1.059 | 0.292 |
| The spatial autocovariate | $1.238 \times 10^{-5}$ | $1.217 \times 10^{-5}$ | 1.017 | 0.312 |
| Response: Soil pH | | | | |
| **Intercept** | 4.414 | $3.226 \times 10^{-1}$ | 13.683 | **<0.001** |
| Land use history | $1.173 \times 10^{-1}$ | $6.614 \times 10^{-2}$ | 1.774 | 0.079 |
| Melanic index | $3.035 \times 10^{-1}$ | $1.678 \times 10^{-1}$ | 1.808 | 0.073 |
| Slope angle | $- 2.890 \times 10^{-3}$ | $4.871 \times 10^{-3}$ | - 0.593 | 0.554 |

| | | | | |
|---|---|---|---|---|
| **The spatial autocovariate** | $3.421 \times 10^{-6}$ | $1.449 \times 10^{-6}$ | 2.361 | **<0.05** |
| Response: Duration of direct sunlight | | | | |
| **Intercept** | $6.858 \times 10^{2}$ | $1.025 \times 10^{2}$ | 6.693 | **<0.001** |
| **Land use history** | $- 4.781 \times 10$ | $2.106 \times 10$ | - 2.270 | **<0.05** |
| Melanic index | $3.872 \times 10$ | $5.326 \times 10$ | 0.727 | 0.469 |
| Slope angle | $- 5.538 \times 10^{-1}$ | 1.540 | - 0.360 | 0.720 |
| The spatial autocovariate | $3.347 \times 10^{-6}$ | $3.128 \times 10^{-6}$ | 1.070 | 0.287 |

| **Predictors** | **Estimate** | **SE** | ***z*** | ***P*** |
|---|---|---|---|---|
| Response: Total species richness | | | | |
| **Intercept** | 2.650 | $3.438 \times 10^{-1}$ | 7.706 | **<0.001** |
| **Land use history** | $- 1.059 \times 10^{-1}$ | $4.070 \times 10^{-2}$ | - 2.602 | **<0.01** |
| **Melanic index** | $- 2.091 \times 10^{-1}$ | $1.013 \times 10^{-1}$ | - 2.065 | **<0.05** |
| Vegetation height | $- 1.058 \times 10^{-3}$ | $5.637 \times 10^{-4}$ | - 1.878 | 0.060 |
| **Aboveground biomass** | $- 4.787 \times 10^{-3}$ | $1.637 \times 10^{-3}$ | - 2.924 | **<0.01** |
| **Soil pH** | $2.836 \times 10^{-1}$ | $6.029 \times 10^{-2}$ | 4.704 | **<0.001** |
| **Duration of direct sunlight** | $- 5.505 \times 10^{-4}$ | $1.801 \times 10^{-4}$ | - 3.058 | **<0.01** |
| **Slope angle** | $5.729 \times 10^{-3}$ | $2.892 \times 10^{-3}$ | 1.981 | **<0.05** |
| **The spatial autocovariate** | $6.597 \times 10^{-7}$ | $9.209 \times 10^{-8}$ | 7.164 | **<0.001** |
| Response: Native grassland species richness | | | | |
| **Intercept** | 2.218 | $4.258 \times 10^{-1}$ | 5.209 | **<0.001** |
| **Land use history** | $- 1.892 \times 10^{-1}$ | $4.911 \times 10^{-2}$ | - 3.853 | **<0.001** |
| **Melanic index** | $- 4.036 \times 10^{-1}$ | $1.272 \times 10^{-1}$ | - 3.172 | **<0.01** |
| Vegetation height | $- 5.904 \times 10^{-4}$ | $6.945 \times 10^{-4}$ | - 0.850 | 0.395 |
| **Aboveground biomass** | $- 4.854 \times 10^{-3}$ | $2.052 \times 10^{-3}$ | - 2.365 | **<0.05** |
| **Soil pH** | $2.935 \times 10^{-1}$ | $7.300 \times 10^{-2}$ | 4.020 | **<0.001** |
| Duration of direct sunlight | $- 7.120 \times 10^{-5}$ | $2.174 \times 10^{-4}$ | - 0.327 | 0.743 |
| Slope angle | $2.187 \times 10^{-3}$ | $3.580 \times 10^{-3}$ | 0.611 | 0.541 |
| **The spatial autocovariate** | $1.164 \times 10^{-6}$ | $1.403 \times 10^{-7}$ | 8.298 | **<0.001** |

| **Predictors** | **Estimate** | **SE** | ***t*** | ***P*** |
|---|---|---|---|---|
| Response: Red list index | | | | |

| | | | | |
|---|---|---|---|---|
| **Intercept** | $- 2.444$ | $9.991 \times 10^{-1}$ | $- 2.446$ | **<0.05** |
| **Land use history** | $- 2.722 \times 10^{-1}$ | $9.512 \times 10^{-2}$ | $- 2.861$ | **<0.01** |
| **Melanic index** | $- 9.321 \times 10^{-1}$ | $3.306 \times 10^{-1}$ | $- 2.820$ | **<0.01** |
| Vegetation height | $- 1.189 \times 10^{-4}$ | $1.648 \times 10^{-3}$ | $0.072$ | 0.943 |
| **Aboveground biomass** | $- 1.574 \times 10^{-2}$ | $5.807 \times 10^{-3}$ | $- 2.711$ | **<0.01** |
| **Soil pH** | $6.876 \times 10^{-1}$ | $1.677 \times 10^{-1}$ | $4.101$ | **<0.001** |
| **Duration of direct sunlight** | $1.075 \times 10^{-3}$ | $4.404 \times 10^{-4}$ | $2.442$ | **<0.05** |
| Slope angle | $- 6.811 \times 10^{-3}$ | $8.080 \times 10^{-3}$ | $- 0.843$ | 0.401 |
| **The spatial autocovariate** | $2.776 \times 10^{-5}$ | $3.766 \times 10^{-6}$ | $7.372$ | **<0.001** |

| **Predictors** | **Estimate** | **SE** | ***z*** | ***P*** |
|---|---|---|---|---|
| Response: Exotic species richness | | | | |
| Intercept | $- 1.992 \times 10^{-1}$ | $1.080$ | $- 0.184$ | 0.854 |
| Land use history | $9.268 \times 10^{-2}$ | $1.318 \times 10^{-1}$ | $0.703$ | 0.482 |
| **Melanic index** | $6.308 \times 10^{-1}$ | $2.799 \times 10^{-1}$ | $2.253$ | **<0.05** |
| Vegetation height | $- 3.228 \times 10^{-3}$ | $1.755 \times 10^{-3}$ | $- 1.839$ | 0.066 |
| Aboveground biomass | $- 4.431 \times 10^{-3}$ | $5.285 \times 10^{-3}$ | $- 0.838$ | 0.402 |
| Soil pH | $1.037 \times 10^{-1}$ | $1.896 \times 10^{-1}$ | $0.547$ | 0.584 |
| **Duration of direct sunlight** | $- 1.375 \times 10^{-3}$ | $5.677 \times 10^{-4}$ | $- 2.421$ | **<0.05** |
| Slope angle | $- 8.465 \times 10^{-4}$ | $8.972 \times 10^{-3}$ | $- 0.094$ | 0.925 |
| **The spatial autocovariate** | $1.936 \times 10^{-5}$ | $2.573 \times 10^{-6}$ | $7.524$ | **<0.001** |
| Response: Endangered grassland species richness | | | | |
| **Intercept** | $- 8.778$ | $4.234$ | $- 2.073$ | **<0.05** |
| **Land use history** | $- 8.474 \times 10^{-1}$ | $3.901 \times 10^{-1}$ | $- 2.172$ | **<0.05** |
| **Melanic index** | $- 3.348$ | $1.495$ | $- 2.240$ | **<0.05** |
| Vegetation height | $5.613 \times 10^{-3}$ | $6.871 \times 10^{-3}$ | $0.817$ | 0.414 |
| Aboveground biomass | $- 1.902 \times 10^{-2}$ | $2.132 \times 10^{-2}$ | $- 0.892$ | 0.372 |
| **Soil pH** | $1.654$ | $6.684 \times 10^{-1}$ | $2.474$ | **<0.05** |
| **Duration of direct sunlight** | $6.226 \times 10^{-3}$ | $2.165 \times 10^{-3}$ | $2.876$ | **<0.01** |
| Slope angle | $2.362 \times 10^{-3}$ | $2.798 \times 10^{-2}$ | $0.084$ | 0.933 |
| **The spatial autocovariate** | $8.308 \times 10^{-5}$ | $2.538 \times 10^{-5}$ | $3.273$ | **<0.01** |
| **Generalized liner mixed model** | | | | |

| **Predictors** | **Estimate** | **SE** | ***z*** | ***P*** |
|---|---|---|---|---|
| Response: Abundances in pasture plots of each species | | | | |
| Intercept | $7.242 \times 10^{-3}$ | $1.562 \times 10^{-2}$ | 0.464 | 0.643 |
| **Abundances in forest plots** | $4.509 \times 10^{-1}$ | $7.401 \times 10^{-2}$ | 6.092 | **<0.001** |
| **Species group (other species including exotic species)** | $6.160 \times 10^{-2}$ | $2.069 \times 10^{-2}$ | 2.978 | **<0.01** |
| **Abundance in forest plots × species group** | $1.943 \times 10^{-1}$ | $9.804 \times 10^{-2}$ | 1.982 | **<0.05** |

**Generalized liner model**

| **Predictors** | **Estimate** | **SE** | ***z*** | ***P*** |
|---|---|---|---|---|
| Response: Barochorous native grassland species richness | | | | |
| **Intercept** | 1.421 | $6.203 \times 10^{-1}$ | 2.290 | **<0.05** |
| **Land use history** | $- 2.074 \times 10^{-1}$ | $6.873 \times 10^{-2}$ | - 3.018 | **<0.01** |
| **Melanic index** | $- 4.881 \times 10^{-1}$ | $1.849 \times 10^{-1}$ | - 2.640 | **<0.01** |
| Vegetation height | $- 1.851 \times 10^{-3}$ | $1.013 \times 10^{-3}$ | - 1.827 | 0.068 |
| **Aboveground biomass** | $- 7.339 \times 10^{-3}$ | $3.060 \times 10^{-3}$ | - 2.398 | **<0.05** |
| **Soil pH** | $3.430 \times 10^{-1}$ | $1.056 \times 10^{-1}$ | 3.249 | **<0.01** |
| Duration of direct sunlight | $4.247 \times 10^{-5}$ | $3.057 \times 10^{-4}$ | 0.139 | 0.890 |
| Slope angle | $- 1.595 \times 10^{-3}$ | $5.194 \times 10^{-3}$ | - 0.307 | 0.759 |
| **The spatial autocovariate** | $2.998 \times 10^{-6}$ | $3.604 \times 10^{-7}$ | 8.317 | **<0.001** |
| Response: Ballochorous native grassland species richness | | | | |
| **Intercept** | - 9.065 | 3.120 | - 2.906 | **<0.01** |
| Land use history | $- 3.320 \times 10^{-1}$ | $3.802 \times 10^{-1}$ | - 0.873 | 0.383 |
| Melanic index | 1.121 | $7.057 \times 10^{-1}$ | 1.589 | 0.112 |
| Vegetation height | $3.273 \times 10^{-3}$ | $4.870 \times 10^{-3}$ | 0.672 | 0.502 |
| Aboveground biomass | $1.030 \times 10^{-2}$ | $1.436 \times 10^{-2}$ | 0.718 | 0.473 |
| Soil pH | $8.056 \times 10^{-1}$ | $5.502 \times 10^{-1}$ | 1.464 | 0.143 |
| Duration of direct sunlight | $6.264 \times 10^{-4}$ | $1.619 \times 10^{-3}$ | 0.387 | 0.699 |
| Slope angle | $- 3.308 \times 10^{-3}$ | $2.363 \times 10^{-2}$ | - 0.140 | 0.889 |
| **The spatial autocovariate** | $1.250 \times 10^{-4}$ | $2.624 \times 10^{-5}$ | 4.761 | **<0.001** |
| Response: Hydrochorous native grassland species richness | | | | |

| | | | | |
|---|---|---|---|---|
| Intercept | - 3.441 | 5.117 | - 0.672 | 0.501 |
| Land use history | $8.185 \times 10^{-1}$ | $7.197 \times 10^{-1}$ | 1.137 | 0.255 |
| Melanic index | $- 5.724 \times 10^{-1}$ | 1.332 | - 0.430 | 0.667 |
| Vegetation height | $- 1.733 \times 10^{-3}$ | $8.406 \times 10^{-3}$ | - 0.206 | 0.837 |
| Aboveground biomass | $- 1.902 \times 10^{-3}$ | $2.361 \times 10^{-2}$ | - 0.081 | 0.936 |
| Soil pH | 1.336 | $9.480 \times 10^{-1}$ | 1.409 | 0.159 |
| Duration of direct sunlight | $- 4.043 \times 10^{-3}$ | $2.520 \times 10^{-3}$ | - 1.605 | 0.109 |
| **Slope angle** | $- 1.406 \times 10^{-1}$ | $4.795 \times 10^{-2}$ | - 2.932 | **<0.01** |
| The spatial autocovariate | $6.060 \times 10^{-5}$ | $2.586 \times 10^{-5}$ | 2.343 | **<0.05** |
| Response: Myrmecochorous native grassland species richness | | | | |
| Intercept | 1.311 | 1.686 | 0.778 | 0.437 |
| Land use history | $- 1.697 \times 10^{-1}$ | $2.004 \times 10^{-1}$ | - 0.847 | 0.397 |
| Melanic index | $- 1.159 \times 10^{-1}$ | $5.159 \times 10^{-1}$ | - 0.225 | 0.822 |
| Vegetation height | $2.153 \times 10^{-5}$ | $2.753 \times 10^{-3}$ | 0.008 | 0.994 |
| Aboveground biomass | $- 7.899 \times 10^{-5}$ | $7.642 \times 10^{-3}$ | - 0.010 | 0.992 |
| Soil pH | $- 2.215 \times 10^{-1}$ | $2.918 \times 10^{-1}$ | - 0.759 | 0.448 |
| Duration of direct sunlight | $- 1.334 \times 10^{-4}$ | $9.122 \times 10^{-4}$ | - 0.146 | 0.884 |
| **Slope angle** | $2.882 \times 10^{-2}$ | $1.416 \times 10^{-2}$ | 2.035 | **<0.05** |
| The spatial autocovariate | $3.062 \times 10^{-6}$ | $9.338 \times 10^{-6}$ | 0.328 | 0.743 |
| Response: Anemochorous native grassland species richness | | | | |
| Intercept | 1.104 | $7.195 \times 10^{-1}$ | 1.535 | 0.125 |
| **Land use history** | $- 2.142 \times 10^{-1}$ | $8.504 \times 10^{-2}$ | - 2.518 | **<0.05** |
| Melanic index | $- 2.848 \times 10^{-1}$ | $2.174 \times 10^{-1}$ | - 1.310 | 0.190 |
| Vegetation height | $5.215 \times 10^{-4}$ | $1.178 \times 10^{-3}$ | 0.443 | 0.658 |
| Aboveground biomass | $- 2.699 \times 10^{-3}$ | $3.445 \times 10^{-3}$ | - 0.784 | 0.433 |
| **Soil pH** | $2.646 \times 10^{-1}$ | $1.240 \times 10^{-1}$ | 2.135 | **<0.05** |
| Duration of direct sunlight | $- 3.140 \times 10^{-4}$ | $3.691 \times 10^{-4}$ | - 0.851 | 0.395 |
| Slope angle | $3.338 \times 10^{-3}$ | $6.140 \times 10^{-3}$ | 0.544 | 0.587 |
| **The spatial autocovariate** | $3.489 \times 10^{-6}$ | $6.638 \times 10^{-7}$ | 5.257 | **<0.001** |
| Response: Epizoochorous native grassland species richness | | | | |
| Intercept | $6.857 \times 10^{-1}$ | 3.129 | 0.219 | 0.827 |

| | | | | |
|---|---|---|---|---|
| Land use history | $5.140 \times 10^{-1}$ | $3.429 \times 10^{-1}$ | 1.499 | 0.134 |
| Melanic index | $- 2.389 \times 10^{-2}$ | $6.755 \times 10^{-1}$ | - 0.035 | 0.972 |
| Vegetation height | $- 5.531 \times 10^{-3}$ | $4.640 \times 10^{-3}$ | - 1.192 | 0.233 |
| Aboveground biomass | $- 2.544 \times 10^{-2}$ | $1.395 \times 10^{-2}$ | - 1.823 | 0.068 |
| Soil pH | $- 2.836 \times 10^{-1}$ | $5.478 \times 10^{-1}$ | - 0.518 | 0.605 |
| Duration of direct sunlight | $7.074 \times 10^{-4}$ | $1.576 \times 10^{-3}$ | 0.449 | 0.654 |
| Slope angle | $- 1.916 \times 10^{-2}$ | $2.305 \times 10^{-2}$ | - 0.831 | 0.406 |
| The spatial autocovariate | $7.277 \times 10^{-5}$ | $2.505 \times 10^{-5}$ | 2.905 | **<0.01** |
| Response: Endozoochorous native grassland species richness | | | | |
| Intercept | 1.014 | 1.934 | 0.524 | 0.600 |
| Land use history | $- 1.310 \times 10^{-1}$ | $2.212 \times 10^{-1}$ | - 0.592 | 0.554 |
| **Melanic index** | - 1.718 | $6.382 \times 10^{-1}$ | - 2.692 | **<0.01** |
| Vegetation height | $4.724 \times 10^{-3}$ | $3.030 \times 10^{-3}$ | 1.559 | 0.119 |
| Aboveground biomass | $- 3.860 \times 10^{-3}$ | $8.717 \times 10^{-3}$ | - 0.443 | 0.658 |
| Soil pH | $3.042 \times 10^{-1}$ | $3.091 \times 10^{-1}$ | 0.984 | 0.325 |
| Duration of direct sunlight | $- 1.401 \times 10^{-4}$ | $1.006 \times 10^{-3}$ | - 0.139 | 0.889 |
| Slope angle | $2.631 \times 10^{-2}$ | $1.586 \times 10^{-2}$ | 1.659 | 0.097 |
| The spatial autocovariate | $5.317 \times 10^{-6}$ | $1.068 \times 10^{-5}$ | 0.498 | 0.619 |

**Generalized liner model & Permutation test**

| **Predictors** | **Estimate** | ***P*** |
|---|---|---|
| Response: Jaccard dissimilarity between plots | | |
| Intercept | $8.116 \times 10^{-1}$ | |
| **Distance between plots** | $8.524 \times 10^{-2}$ | **<0.001** |
| **Bray-Curtis dissimilarity** | $1.797 \times 10^{-1}$ | **<0.001** |
| **Land use history** | | |
| **Forest – forest pair** | $1.079 \times 10^{-2}$ | **<0.001** |
| **Pasture – forest pair** | $7.185 \times 10^{-3}$ | **<0.001** |
| **Distance × f – f pair** | $- 1.712 \times 10^{-2}$ | **<0.001** |
| Distance × p – f pair | $1.603 \times 10^{-3}$ | 0.128 |

| | | |
|---|---|---|
| Dissimilarity × f – f pair | $- 1.948 \times 10^{-2}$ | 1.000 |
| Dissimilarity × p – f pair | $- 5.922 \times 10^{-3}$ | 0.655 |

Table S4. Information on species detected in the study plots: origin (native vs. exotic, N/E); habitat (native grassland species vs. forest species, G/F); dispersal mode (native grassland species only); number of plots in which species occurred on pasture/forest ski slopes, P/F.

| Family | Species | Origin | Habitat | Dispersal | Red list category Nat/Nag/Gum | RLI | No. of plots Pasture | No. of plots Forest |
|---|---|---|---|---|---|---|---|---|
| *Pinaceae* | | | | | | | | |
| | *Pinus densiflora* | N | F | | | 0 | 10 | 0 |
| | *Larix kaempferi* | N | F | | | 0 | 2 | 18 |
| *Schisandraceae* | | | | | | | | |
| | *Schisandra chinensis* | N | F | | | 0.2535 | 0 | 5 |
| *Chloranthaceae* | | | | | | | | |
| | *Chloranthus serratus* | N | F | | | 0 | 0 | 1 |
| *Ranunculaceae* | | | | | | | | |
| | *Aquilegia buergeriana* | N | G | Barochory | | 0.1467 | 5 | 12 |
| | *Pulsatilla cernua* | N | G | Anemochory | VU/VU/EN | 0.7167 | 2 | 0 |
| | *Ranunculus japonicus* | N | G | Barochory | | 0.0106 | 6 | 1 |
| | *Ranunculus silerifolius* | N | G | Barochory | | 0 | 2 | 0 |
| | *Thalictrum aquilegiifolium* | N | G | Barochory | | 0.0714 | 5 | 7 |
| | *Thalictrum minus* | N | G | Barochory | | 0.0217 | 10 | 6 |
| | *Aconitum japonicum* | N | F | | | 0.0500 | 0 | 2 |
| | *Thalictrum tuberiferum* | N | F | | | 0.1196 | 1 | 6 |
| *Lardizabalaceae* | | | | | | | | |
| | *Akebia trifoliata* | N | F | | | 0 | 1 | 0 |
| *Polygonaceae* | | | | | | | | |
| | *Fallopia japonica* | N | G | Anemochory | | 0 | 10 | 31 |

| | | | | | | | |
|---|---|---|---|---|---|---|---|
| *Persicaria lapathifolia* | N | G | Barochory | | 0 | 0 | 1 |
| *Persicaria longiseta* | N | G | Myrmecochory | | 0 | 0 | 2 |
| *Persicaria nepalensis* | N | G | Barochory | | 0.0272 | 0 | 1 |
| *Persicaria posumbu* | N | G | Barochory | | 0.0053 | 0 | 1 |
| *Persicaria sagittata* | N | G | Hydrochory | | 0 | 1 | 3 |
| *Rumex acetosa* | N | G | Anemochory | | 0 | 4 | 0 |
| *Rumex japonicus* | N | G | Anemochory | | 0 | 0 | 3 |
| *Rumex acetosella* | E | G | | | 0 | 8 | 9 |
| *Caryophyllaceae* | | | | | | | |
| *Arenaria lateriflora* | N | G | Myrmecochory | | 0.1739 | 25 | 27 |
| *Dianthus superbus* | N | G | Barochory | | 0.0722 | 1 | 1 |
| *Silene baccifera* | N | G | Endozoochory | | 0.0598 | 0 | 1 |
| *Silene miqueliana* | N | F | | | 0.1576 | 1 | 0 |
| *Cerastium glomeratum* | E | G | | | 0 | 0 | 1 |
| *Santalaceae* | | | | | | | |
| *Thesium chinense* | N | G | Barochory | | 0 | 4 | 0 |
| *Paeoniaceae* | | | | | | | |
| *Paeonia obovata* | N | F | | VU/EN/EN | 0.5326 | 2 | 1 |
| *Saxifragaceae* | | | | | | | |
| *Astilbe microphylla* | N | G | Barochory | | 0.1033 | 6 | 18 |
| *Astilbe odontophylla* | N | G | Barochory | | 0.0317 | 3 | 2 |
| *Astilbe thunbergii* | N | F | | | 0 | 0 | 5 |
| *Rodgersia podophylla* | N | F | | | 0.100 | 4 | 5 |
| *Grossulariaceae* | | | | | | | |
| *Ribes fasciculatum* | N | F | | | 0.1833 | 0 | 1 |
| *Haloragaceae* | | | | | | | |
| *Haloragis micrantha* | N | G | Barochory | | 0.0160 | 1 | 0 |

| | | | | | | |
|---|---|---|---|---|---|---|
| *Malvaceae* | | | | | | |
| *Tilia japonica* | N | F | | 0.0952 | 0 | 1 |
| *Salicaceae* | | | | | | |
| *Populus tremula* | N | F | | 0 | 10 | 2 |
| *Salix caprea* | N | F | | 0.0809 | 8 | 19 |
| *Salix chaenomeloides* | N | F | | 0.0179 | 3 | 12 |
| *Salix integra* | N | F | | 0.0217 | 5 | 11 |
| *Salix pierotii* | N | F | | 0 | 0 | 1 |
| *Salix udensis* | N | F | | 0.0128 | 0 | 6 |
| *Violaceae* | | | | | | |
| *Viola grypoceras* | N | G | Myrmecochory | 0 | 8 | 18 |
| *Viola mandshurica* | N | G | Myrmecochory | 0 | 12 | 0 |
| *Viola phalacrocarpa* | N | G | Myrmecochory | 0.0652 | 4 | 1 |
| *Viola verecunda A.Gray var. semilunaris Maxim.* | N | G | Myrmecochory | 0 | 0 | 1 |
| *Hypericaceae* | | | | | | |
| *Hypericum erectum* | N | G | Barochory | 0 | 25 | 21 |
| *Celastraceae* | | | | | | |
| *Parnassia palustris* | N | G | Barochory | 0.2011 | 5 | 1 |
| *Celastrus orbiculatus* | N | F | | 0 | 18 | 17 |
| *Euonymus alatus* | N | F | | 0 | 5 | 10 |
| *Euonymus sieboldianus* | N | F | | 0 | 0 | 2 |
| *Rosaceae* | | | | | | |
| *Agrimonia pilosa* | N | G | Epizoochory | 0.0163 | 4 | 7 |
| *Filipendula camtschatica* | N | G | Barochory | 0.0326 | 0 | 2 |
| *Filipendula multijuga* | N | G | Barochory | 0.1901 | 0 | 1 |
| *Potentilla cryptotaeniae* | N | G | Barochory | 0.1087 | 1 | 0 |

| | | | | | | |
|---|---|---|---|---|---|---|
| *Potentilla fragarioides* | N | G | Barochory | 0.0163 | 13 | 5 |
| *Potentilla freyniana* | N | G | Barochory | 0 | 43 | 41 |
| *Potentilla hebiichigo* | N | G | Endozoochory | 0 | 0 | 1 |
| *Sanguisorba officinalis* | N | G | Barochory | 0.0543 | 38 | 4 |
| *Aruncus dioicus* | N | F | | 0.0978 | 0 | 3 |
| *Geum japonicum* | N | F | | 0.0054 | 0 | 3 |
| *Kerria japonica* | N | F | | 0.0217 | 0 | 1 |
| *Malus toringo* | N | F | | 0.1630 | 3 | 6 |
| *Rosa multiflora* | N | G | Endozoochory | 0 | 3 | 13 |
| *Rubus minusculus* | N | F | | 0.0634 | 0 | 2 |
| *Rubus parvifolius* | N | G | Endozoochory | 0 | 4 | 10 |
| *Rubus pungens* | N | F | | 0.1889 | 0 | 6 |
| *Rubus subcrataegifolius* | N | G | Endozoochory | 0.0086 | 32 | 40 |
| *Sorbus commixta* | N | F | | 0.0109 | 0 | 3 |
| *Urticaceae* | | | | | | |
| *Boehmeria nivea* | N | G | Barochory | 0 | 0 | 2 |
| *Boehmeria spicata* | N | G | Barochory | 0 | 0 | 1 |
| *Laportea bulbifera* | N | F | | 0.0054 | 0 | 1 |
| *Cannabaceae* | | | | | | |
| *Humulus lupulus* | N | F | | 0.1250 | 4 | 3 |
| *Moraceae* | | | | | | |
| *Morus australis* | N | F | | 0 | 0 | 1 |
| *Geraniaceae* | | | | | | |
| *Geranium thunbergii* | N | G | Ballistic | 0 | 4 | 16 |
| *Geranium yesoense* | N | G | Barochory | 0 | 0 | 1 |
| *Vitaceae* | | | | | | |
| *Vitis coignetiae* | N | F | | 0.0217 | 4 | 3 |
| *Onagraceae* | | | | | | |

| | | | | | | | |
|---|---|---|---|---|---|---|---|
| | *Chamerion angustifolium* | N | G | Anemochory | | 0.2813 | 8 | 13 |
| | *Epilobium amurense* | N | G | Anemochory | | 0.0761 | 0 | 1 |
| | *Circaea erubescens* | N | F | | | 0.0435 | 1 | 3 |
| | *Oenothera biennis* | E | G | | | 0 | 21 | 30 |
| *Betulaceae* | | | | | | | | |
| | *Alnus firma* | N | F | | | 0.0521 | 0 | 1 |
| | *Betula platyphylla* | N | F | | | 0.0313 | 12 | 17 |
| | *Corylus sieboldiana* | N | F | | | 0.0707 | 1 | 3 |
| *Fagaceae* | | | | | | | | |
| | *Castanea crenata* | N | F | | | 0 | 3 | 1 |
| | *Quercus aliena* | N | F | | LC/VU/LC | 0.0739 | 0 | 1 |
| | *Quercus crispula* | N | F | | | 0.0217 | 13 | 9 |
| *Fabaceae* | | | | | | | | |
| | *Amphicarpaea bracteata* | N | G | Ballistic | | 0 | 2 | 14 |
| | *Lespedeza bicolor* | N | G | Barochory | | 0 | 37 | 18 |
| | *Lotus corniculatus* | N | G | Barochory | | 0 | 6 | 3 |
| | *Trifolium lupinaster* | N | G | Barochory | | 0.1875 | 0 | 4 |
| | *Kummerowia striata* | N | F | | | 0 | 0 | 1 |
| | *Trifolium pratense* | E | G | | | 0 | 3 | 16 |
| | *Trifolium repens* | E | G | | | 0 | 5 | 11 |
| *Polygalaceae* | | | | | | | | |
| | *Polygala japonica* | N | G | Myrmecochory | | 0.0106 | 7 | 0 |
| *Anacardiaceae* | | | | | | | | |
| | *Toxicodendron orientale* | N | F | | | 0.0054 | 0 | 2 |
| | *Toxicodendron trichocarpum* | N | F | | | 0 | 0 | 2 |
| *Sapindaceae* | | | | | | | | |

| | | | | | | | |
|---|---|---|---|---|---|---|---|
| *Acer crataegifolium* | N | | | | 0.0125 | 0 | 1 |
| *Acer distylum* | N | F | | | 0.0306 | 0 | 1 |
| *Acer ginnala* | N | F | | | 0.1033 | 4 | 1 |
| *Acer rufinerve* | N | F | | | 0 | 0 | 1 |
| *Acer tschonoskii* | N | F | | | 0 | 0 | 1 |
| *Brassicaceae* | | | | | | | |
| *Arabis hirsuta* | N | G | Barochory | | 0.0109 | 1 | 0 |
| *Barbarea vulgaris* | E | G | | | 0 | 2 | 7 |
| *Sisymbrium luteum* | N | F | | | 0.1890 | 0 | 2 |
| *Turritis glabra* | N | G | Barochory | LC/LC/VU | 0.1522 | 1 | 0 |
| *Asteraceae* | | | | | | | |
| *Achillea alpina* | N | G | Barochory | | 0.1643 | 4 | 0 |
| *Ambrosia artemisiifolia* | E | G | | | 0 | 0 | 1 |
| *Ambrosia trifida* | E | G | | | 0 | 2 | 5 |
| *Anaphalis margaritacea* | N | G | Anemochory | | 0.1071 | 17 | 15 |
| *Artemisia indica* | N | G | Barochory | | 0 | 22 | 21 |
| *Artemisia japonica* | N | G | Barochory | | 0 | 17 | 3 |
| *Artemisia montana* | N | F | | | 0 | 19 | 46 |
| *Aster glehnii* | N | G | Anemochory | | 0.0368 | 5 | 27 |
| *Aster iinumae* | N | G | Barochory | | 0 | 0 | 2 |
| *Aster microcephalus* | N | G | Anemochory | | 0.0056 | 19 | 15 |
| *Aster scaber* | N | G | Anemochory | | 0 | 4 | 2 |
| *Aster semiamplexicaulis* | N | G | Anemochory | | 0 | 1 | 0 |
| *Aster yomena* | N | G | Barochory | | 0 | 2 | 3 |
| *Cirsium japonicum* | N | G | Anemochory | | 0 | 16 | 28 |
| *Cirsium tonense* | N | G | | | 0 | 1 | 5 |
| *Cirsium oligophyllum* | N | G | Anemochory | | 0 | 33 | 8 |
| *Erigeron annuus* | E | G | | | 0 | 10 | 21 |

| | | | | | | | |
|---|---|---|---|---|---|---|---|
| *Erigeron strigosus* | E | G | | | 0 | 24 | 19 |
| *Erigeron thunbergii* | N | G | Anemochory | LC/VU/EN | 0.6167 | 14 | 0 |
| *Eupatorium glehnii* | N | F | | | 0.0656 | 9 | 24 |
| *Eupatorium makinoi* | N | F | | | 0 | 0 | 1 |
| *Hieracium umbellatum* | N | G | Anemochory | | 0.2885 | 6 | 2 |
| *Hypochaeris radicata* | E | G | | | 0 | 4 | 2 |
| *Inula salicina* | N | G | Anemochory | | 0.2554 | 10 | 1 |
| *Ixeridium dentatum* | N | G | Anemochory | | 0 | 29 | 10 |
| *Lactuca indica* | N | G | Anemochory | | 0 | 4 | 0 |
| *Leucanthemum vulgare* | E | G | | | 0 | 0 | 10 |
| *Ligularia dentata* | N | G | Anemochory | | 0.1053 | 7 | 4 |
| *Petasites japonicus* | N | F | | | 0 | 13 | 6 |
| *Picris hieracioides* | N | G | Anemochory | | 0 | 17 | 14 |
| *Rudbeckia laciniata* | E | G | | | 0 | 0 | 1 |
| *Saussurea ussuriensis* | N | G | Anemochory | LC/LC/EN | 0.4653 | 3 | 0 |
| *Senecio cannabifolius* | N | G | Anemochory | | 0.0781 | 8 | 18 |
| *Senecio nemorensis* | N | F | | | 0.1033 | 6 | 5 |
| *Serratula coronata* | N | G | Anemochory | | 0.1167 | 4 | 1 |
| *Solidago virgaurea* | N | G | Anemochory | | 0 | 38 | 23 |
| *Synurus excelsus* | N | G | Anemochory | | 0.1625 | 5 | 1 |
| *Synurus pungens* | N | G | Anemochory | | 0.1100 | 2 | 0 |
| *Taraxacum officinale* | E | G | | | 0 | 4 | 4 |
| *Tephroseris flammea subsp. glabrifolia* | N | G | Anemochory | VU/LC/VU | 0.2647 | 3 | 0 |
| *Tephroseris integrifolia* | N | G | Anemochory | | 0.2111 | 5 | 0 |
| *Campanulaceae* | | | | | | | |

| | | | | | | | |
|---|---|---|---|---|---|---|---|
| *Adenophora triphylla* | N | G | Barochory | | 0.0109 | 25 | 4 |
| *Campanula punctata* | N | F | | | 0.0778 | 11 | 4 |
| *Platycodon grandiflorus* | N | G | Barochory | VU/NT/VU | 0.4783 | 19 | 0 |
| *Lamiaceae* | | | | | | | |
| *Agastache rugosa* | N | G | Barochory | | 0.1359 | 0 | 2 |
| *Chelonopsis moschata* | N | F | | | 0.1141 | 1 | 0 |
| *Clinopodium chinense* | N | G | Barochory | | 0 | 3 | 7 |
| *Clinopodium micranthum* | N | F | | | 0 | 4 | 1 |
| *Lycopus maackianus* | N | G | Barochory | | 0.0707 | 0 | 2 |
| *Prunella vulgaris* | N | G | Barochory | | 0 | 32 | 11 |
| *Stachys aspera* | N | G | Barochory | | 0.0707 | 0 | 1 |
| *Teucrium japonicum* | N | G | Barochory | | 0.0106 | 0 | 1 |
| *Thymus quinquecostatus* | N | G | Anemochory | | 0.1250 | 23 | 1 |
| *Plantaginaceae* | | | | | | | |
| *Plantago asiatica* | N | G | Epizoochory | | 0 | 6 | 24 |
| *Plantago lanceolata* | E | G | | | 0 | 6 | 2 |
| *Veronica onoei* | N | G | Anemochory | VU/NT/VU | 0.1389 | 8 | 0 |
| *Veronicastrum japonicum* | N | G | Barochory | | 0.1029 | 3 | 10 |
| *Oleaceae* | | | | | | | |
| *Fraxinus lanuginosa* | N | F | | | 0.0163 | 0 | 1 |
| *Ligustrum tschonoskii* | N | F | | | 0.0489 | 1 | 3 |
| *Orobanchaceae* | | | | | | | |
| *Euphrasia maximowiczii* | N | G | Barochory | | 0.2864 | 10 | 6 |
| *Apiaceae* | | | | | | | |

| | | | | | | | |
|---|---|---|---|---|---|---|---|
| *Angelica decursiva* | N | G | Anemochory | | 0.0214 | 0 | 2 |
| *Angelica pubescens* | N | G | Anemochory | | 0.0056 | 12 | 18 |
| *Cryptotaenia canadensis* | N | F | | | 0 | 2 | 8 |
| *Hydrocotyle maritima* | N | G | Barochory | | 0 | 1 | 0 |
| *Hydrocotyle ramiflora* | N | G | Barochory | | 0 | 18 | 1 |
| *Libanotis ugoensis* | N | G | Barochory | | 0.2167 | 9 | 6 |
| *Araliaceae* | | | | | | | |
| *Aralia cordata* | N | G | Endozoochory | | 0.0054 | 4 | 22 |
| *Aralia elata* | N | F | | | 0 | 0 | 4 |
| *Balsaminaceae* | | | | | | | |
| *Impatiens noli-tangere* | N | G | Ballistic | | 0.0543 | 1 | 7 |
| *Ericaceae* | | | | | | | |
| *Gaultheria pyroloides* | N | G | Endozoochory | | 0.1176 | 0 | 3 |
| *Pyrola asarifolia* | N | F | | | 0.2656 | 1 | 0 |
| *Rhododendron molle* | N | F | | | 0.2011 | 4 | 1 |
| *Primulaceae* | | | | | | | |
| *Lysimachia clethroides* | N | G | Barochory | | 0 | 32 | 15 |
| *Lysimachia japonica* | N | G | Barochory | | 0 | 7 | 7 |
| *Trientalis europaea* | N | G | Barochory | | 0.1026 | 0 | 2 |
| *Symplocaceae* | | | | | | | |
| *Symplocos sawafutagi* | N | F | | | 0 | 1 | 0.0054 |
| *Caprifoliaceae* | | | | | | | |
| *Patrinia scabiosifolia* | N | G | Barochory | LC/LC/VU | 0.1685 | 12 | 0 |
| *Patrinia villosa* | N | G | Barochory | | 0.0109 | 0 | 3 |
| *Scabiosa japonica* | N | G | Barochory | | 0.4076 | 14 | 0 |
| *Triosteum sinuatum* | N | G | Barochory | VU/VU/LC | 0.5 | 2 | 0 |

| | | | | | | | |
|---|---|---|---|---|---|---|---|
| *Weigela decora* | N | F | | | 0.0316 | 0 | 2 |
| *Adoxaceae* | | | | | | | |
| *Sambucus racemosa* | N | F | | | 0 | 1 | 1 |
| *Cornaceae* | | | | | | | |
| *Cornus controversa* | N | F | | | 0 | 0 | 2 |
| *Cornus macrophylla* | N | F | | | 0 | 0 | 2 |
| *Hydrangeaceae* | | | | | | | |
| *Hydrangea paniculata* | N | F | | | 0.0163 | 0 | 18 |
| *Boraginaceae* | | | | | | | |
| *Cynoglossum asperrimum* | N | F | | | 0.0815 | 0 | 1 |
| *Lithospermum erythrorhizon* | N | G | Barochory | EN/EN/EN | 0.5924 | 1 | 0 |
| *Apocynaceae* | | | | | | | |
| *Cynanchum caudatum* | N | G | Anemochory | | 0.0543 | 3 | 4 |
| *Vincetoxicum atratum* | N | G | Anemochory | VU/VU/EN | 0.5435 | 2 | 0 |
| *Vincetoxicum pycnostelma* | N | G | Anemochory | NT/NT/EN | 0.4348 | 3 | 0 |
| *Gentianaceae* | | | | | | | |
| *Gentiana scabra* | N | G | Barochory | | 0.0778 | 13 | 1 |
| *Gentiana triflora* | N | G | Barochory | | 0.1406 | 5 | 2 |
| *Gentiana zollingeri* | N | G | Hydrochory | | 0.0326 | 2 | 1 |
| *Halenia corniculata* | N | G | Hydrochory | | 0.1304 | 1 | 0 |
| *Tripterospermum japonicum* | N | F | | | 0.0326 | 0 | 1 |
| *Rubiaceae* | | | | | | | |
| *Galium japonicum* | N | F | | | 0.0380 | 2 | 3 |
| *Galium trachyspermum* | N | G | Barochory | | 0.0109 | 0 | 4 |
| *Galium verum* | N | G | Barochory | | 0.0489 | 38 | 5 |

| | | | | | | | |
|---|---|---|---|---|---|---|---|
| *Araceae* | | | | | | | |
| *Arisaema japonicum* | N | G | Endozoochory | | 0 | 0 | 1 |
| *Xanthorrhoeaceae* | | | | | | | |
| *Hemerocallis citrina* | N | G | Barochory | LC/NT/LC | 0.2333 | 2 | 1 |
| *Asparagaceae* | | | | | | | |
| *Convallaria majalis* | N | G | Endozoochory | | 0.1786 | 30 | 13 |
| *Hosta sieboldiana* | N | G | Anemochory | | 0.0380 | 23 | 8 |
| *Maianthemum dilatatum* | N | F | | | 0.0761 | 0 | 5 |
| *Polygonatum humile* | N | F | | LC/LC/EN | 0.1467 | 0 | 1 |
| *Polygonatum odoratum* | N | G | Endozoochory | | 0.0109 | 20 | 5 |
| *Iridaceae* | | | | | | | |
| *Iris sanguinea* | N | G | Barochory | | 0.1793 | 38 | 14 |
| *Orchidaceae* | | | | | | | |
| *Liparis japonica* | N | F | | LC/EN/EN | 0.4402 | 1 | 0 |
| *Platanthera sachalinensis* | N | F | | | 0.2283 | 0 | 2 |
| *Spiranthes sinensis* | N | G | Anemochory | | 0 | 0 | 3 |
| *Dioscoreaceae* | | | | | | | |
| *Dioscorea nipponica* | N | F | | | 0.1250 | 7 | 13 |
| *Nartheciaceae* | | | | | | | |
| *Aletris foliata* | N | G | Epizoochory | | 0.1204 | 4 | 11 |
| *Metanarthecium luteoviride* | N | F | | | 0.0217 | 0 | 2 |
| *Colchicaceae* | | | | | | | |
| *Disporum sessile* | N | G | Endozoochory | | 0.0109 | 0 | 3 |
| *Disporum smilacinum* | N | G | Endozoochory | | 0.0435 | 4 | 6 |
| *Liliaceae* | | | | | | | |
| *Lilium leichtlinii* | N | G | Barochory | | 0.0489 | 7 | 2 |
| *Lilium medeoloides* | N | G | Anemochory | | 0.1048 | 0 | 1 |

| | | | | | | | | |
|---|---|---|---|---|---|---|---|---|
| *Melanthiaceae* | | | | | | | | |
| | *Helonias orientalis* | N | G | Barochory | | 0 | 1 | 1 |
| *Poaceae* | | | | | | | | |
| | *Anthoxanthum odoratum* | E | G | | | 0 | 11 | 2 |
| | *Arthraxon hispidus* | N | G | Barochory | | 0 | 0 | 7 |
| | *Arundinella hirta* | N | G | Barochory | | 0 | 19 | 5 |
| | *Bromus remotiflorus* | N | G | Barochory | | 0 | 0 | 11 |
| | *Dactylis glomerata* | E | G | | | 0 | 12 | 30 |
| | *Elymus tsukushiensis* | N | G | Epizoochory | | 0 | 0 | 5 |
| | *Festuca extremiorientalis* | N | F | | | 0.1094 | 0 | 1 |
| | *Festuca ovina* | N | G | Barochory | LC/LC/VU | 0.0833 | 18 | 10 |
| | *Festuca rubra* | E | G | | | 0 | 3 | 1 |
| | *Lolium arundinaceum* | E | G | | | 0 | 10 | 5 |
| | *Lolium pratense* | E | G | | | 0 | 1 | 2 |
| | *Imperata cylindrica* | N | G | Anemochory | | 0 | 1 | 0 |
| | *Miscanthus sinensis* | N | G | Anemochory | | 0 | 43 | 23 |
| | *Phleum pratense* | E | G | | | 0 | 3 | 26 |
| | *Poa sphondylodes* | N | G | Barochory | | 0.0163 | 13 | 40 |
| | *Poa trivialis* | E | G | | | 0 | 0 | 1 |
| | *Sasa senanensis* | N | F | | | 0 | 22 | 15 |
| | *Scirpus wichurae* | N | G | Hydrochory | | 0 | 0 | 2 |
| | *Spodiopogon sibiricus* | N | G | Barochory | | 0.038 | 2 | 1 |
| | *Zoysia japonica* | N | G | Barochory | | 0 | 22 | 4 |
| | *sp 1* | | | | | | 1 | 0 |
| | *sp 2* | | | | | | 0 | 4 |
| | *sp 3* | | | | | | 0 | 1 |
| *Cyperaceae* | | | | | | | | |
| | *Carex dickinsii* | N | G | Hydrochory | | 0.1196 | 0 | 1 |

| | | | | | | | | |
|---|---|---|---|---|---|---|---|---|
| | *Carex dispalata* | N | G | Hydrochory | | 0.0163 | 1 | 0 |
| | *Carex fernaldiana* | N | F | | | 0.0598 | 0 | 3 |
| | *Carex hakonensis* | N | F | | | 0.1141 | 0 | 1 |
| | *Carex incisa* | N | G | Barochory | | 0 | 1 | 0 |
| | *Carex lanceolata* | N | G | Barochory | | 0.0163 | 34 | 19 |
| | *Carex leucochlora* | N | G | Myrmecochory | | 0 | 7 | 6 |
| | *Carex maximowiczii* | N | G | Hydrochory | | 0.0053 | 0 | 2 |
| | *Carex nervata* | N | G | Barochory | | 0.0272 | 23 | 18 |
| | *Carex oxyandra* | N | G | Barochory | | 0.1196 | 0 | 1 |
| | *Carex thunbergii* | N | G | Barochory | | 0.0857 | 0 | 1 |
| | *Fimbristylis complanata* | N | G | Barochory | LC/LC/VU | 0.1250 | 0 | 2 |
| | *sp 4* | | | | | | 0 | 2 |
| *Juncaceae* | | | | | | | | |
| | *Juncus decipiens* | N | G | Hydrochory | | 0.0160 | 0 | 5 |
| | *Luzula capitata* | N | G | Myrmecochory | | 0.0160 | 3 | 1 |
| | *Luzula multiflora* | N | G | Myrmecochory | | 0.0385 | 6 | 10 |
| *Lycopodiaceae* | | | | | | | | |
| | *Lycopodium clavatum* | N | G | Anemochory | | 0.0326 | 1 | 1 |
| *Athyriaceae* | | | | | | | | |
| | *Athyrium yokoscense* | N | G | Anemochory | | 0.0054 | 0 | 2 |
| *Dennstaedtiaceae* | | | | | | | | |
| | *Pteridium aquilinum* | N | G | Anemochory | | 0 | 34 | 11 |
| *Dryopteridaceae* | | | | | | | | |
| | *Arachniodes rhomboidea* | N | F | | | 0.0813 | 0 | 1 |
| | *Dryopteris crassirhizoma* | N | F | | | 0.0705 | 0 | 1 |
| | *Polystichum polyblepharon* | N | F | | | 0 | 0 | 2 |
| *Thelypteridaceae* | | | | | | | | |

| | | | | | | |
|---|---|---|---|---|---|---|
| | *Thelypteris bukoensis* | N | F | | 0.7045 | 0 | 1 |
| | *Thelypteris glanduligera* | N | F | | 0.0427 | 0 | 4 |
| | *Thelypteris palustris* | N | G | Anemochory | 0.0326 | 0 | 1 |
| | *Thelypteris torresiana* | N | G | Anemochory | 0.0227 | 2 | 2 |
| | *sp 5* | | | | | 0 | 6 |
| *Osmundaceae* | | | | | | | |
| | *Osmunda claytoniana* | N | G | Anemochory | 0.1290 | 0 | 5 |
| | *Osmunda japonica* | N | F | | 0.0213 | 0 | 1 |
| *Equisetaceae* | | | | | | | |
| | *Equisetum arvense* | N | G | Anemochory | 0 | 3 | 2 |
| *Ophioglossaceae* | | | | | | | |
| | *Botrychium ternatum* | N | G | Anemochory | 0.0372 | 4 | 6 |
| | *Botrychium virginianum* | N | F | | 0.1059 | 0 | 1 |
| | *Ophioglossum vulgatum* | N | G | Anemochory | 0.2717 | 0 | 3 |
